# Effector loss and gain drives pathogen host range at a fitness cost

**DOI:** 10.1101/2025.09.25.678615

**Authors:** Marcus V. Merfa, Tracy E. Hawk, Jelmer W. Poelstra, Lillian Ebeling-Koning, Ervan Rodgers, Hannah Toth, Zachary Konkel, Jules Butchacas, Nathaniel Heiden, Muhammad Uzair, Temitope F. Oladele, Godwin E. Oduokpaha, Rebecca D. Curland, Emmanuelle Lauber, Laurie Marsan, Zhaohui Liu, Ruth Dill-Macky, Laurent D. Noël, Horacio D. Lopez-Nicora, Jason C. Slot, Verónica Román-Reyna, Jonathan M. Jacobs

## Abstract

Epidemic preparedness depends on tracking microbial evolution that drives shifts in ecological behaviors such as disease emergence. However, the genetic constraints for microbial host adaptation to emerge for generalist and specialist behaviors remain poorly described. Here, we show that generalist cereal pathogen *Xanthomonas translucens* arose from a specialist ancestor via the loss of a single effector gene, *xopAL1*. Deleting barley-specialist *X. translucens xopAL1* recapitulated the host jump to wheat and demonstrates risk across each globally distributed genetic lineage. However, this niche expansion via XopAL1 loss incurs a significant pathogenic fitness cost to colonize barley. Moreover, the specialist lineage gained an additional effector gene, *xopAJ*, which enhanced virulence on barley while restricting oat infection, thereby reinforcing niche specialization. We further identified key host pathways mediating resistance to the specialist lineage of *X. translucens*, opening avenues for potentially identifying targets for crop improvement. Our work provides an experimentally validated evolutionary framework to understand mechanisms of intergenera host jump. Overall, we demonstrate that single events of gene loss and gain shape ecological behaviors of pathogens by creating a dynamic trade-off between niche breadth and specialization.

## Introduction

Disease outbreaks often result from pathogen evolution, leading to changes in microbial ecological behaviors such as niche specialization. Changes in niche-specificity may occur from single genetic loss or gain events^1,2^ or the movement of genomic islands and plasmids that dictate the transitional behaviors such as niche shift between microbial pathogens and their hosts^3^. Niche alterations are universal and widespread in plant, human and animal epidemiology and have significant effect on host health outcomes^4^. Some pathogens have generalist behaviors with broad host ranges while others are highly specialized on a particular host or niche^5^. However, the underlying evolutionary and biological mechanisms often remain undefined for most niche changes, especially in plant agricultural systems.

Host range is a particular type of niche-specificity, and many pathogenic organisms can be found across the spectrum from generalist to specialist. The Gammaproteobacterial genus *Xanthomonas* limits agricultural production by causing diseases in over 400 identified plant species, although these pathogens as a whole infect close to 50 plant families. *Xanthomonas* bacteria in the generalist spectrum cause disease in a broad range of host genera with access to a breadth of potential hosts for resources, while specialists remain restricted to a particular species or genus (Fig. 1A and Fig. S1)^6–8^. Curiously, differences in niche-specificity of closely related plant pathogenic species and strains are often due to small changes in genome content^9–12^ but the evolutionary mechanisms for niche specialization of *Xanthomonas* spp. remains poorly understood. Ultimately, understanding the spectrum and evolution of host range specificity will benefit prediction and mitigation of disease epidemics^5,13^.

**Fig. 1.**
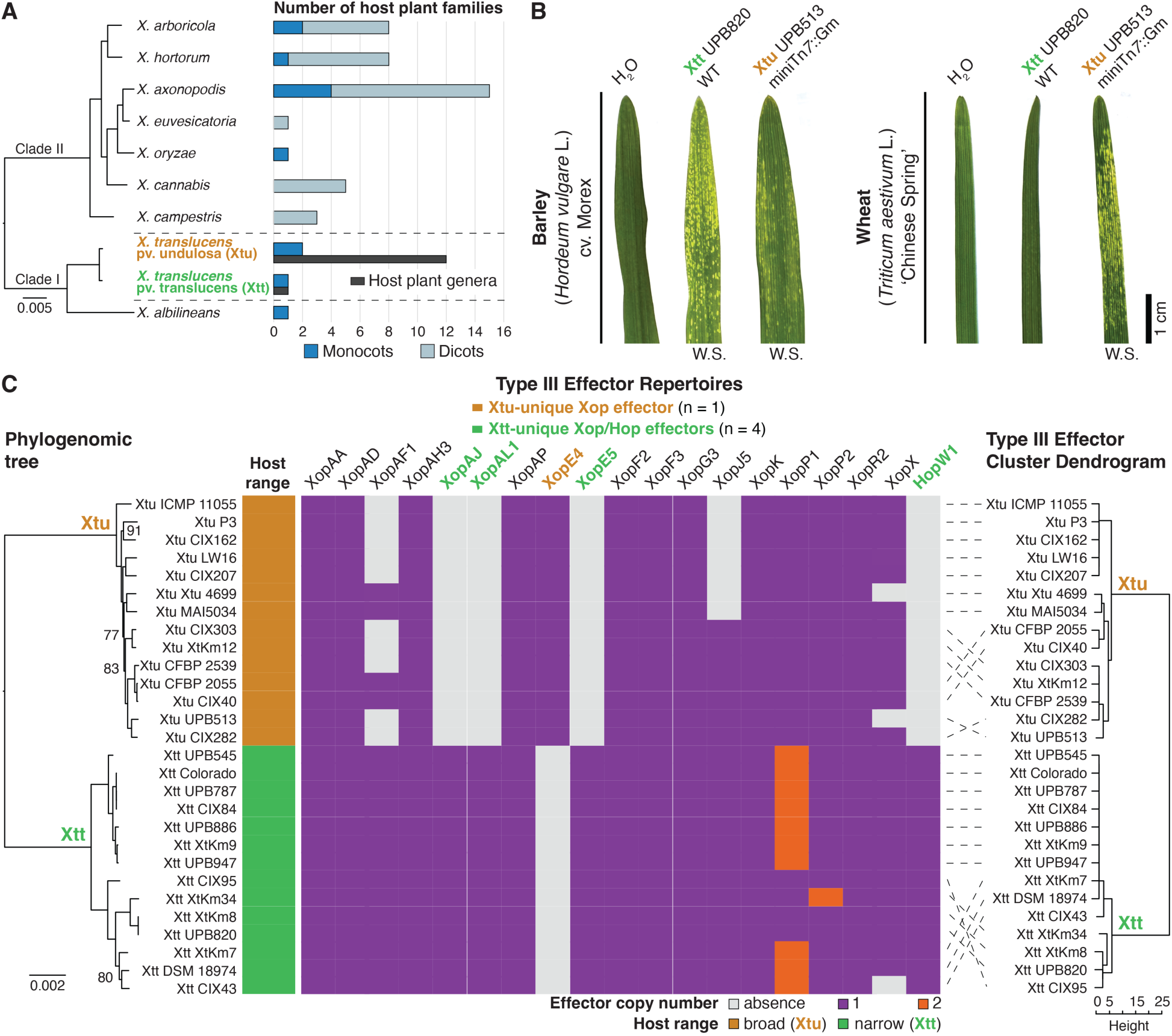
*X. translucens* pathovars with differential host range breadth encode significantly different effector repertoires. **A.** Phylogenomics and host range of representative *Xanthomonas* spp. The number of plant families that are known hosts (natural or experimentally determined) for each analyzed *Xanthomonas* spp. are shown in the chart to the right side of the panel. The number of plant genera that are known hosts for *X. translucens* pathovars translucens and undulosa is also included in the chart. **B.** Symptom development in barley (*Hordeum vulgare* L. cv. Morex) and wheat (*Triticum aestivum* L. genotype ‘Chinese Spring’) leaves inoculated (spray) with Xtu and Xtt strains (OD_600_ 0.5) at six days post inoculation. Water soaking symptoms were visualized using a light box and appear as lighter areas in the leaf. Xtt elicited no response in wheat. Scale bar is shown in figure. W.S. – water soaking. **C.** Effector repertoires of *X. translucens* pvs. translucens and undulosa (only non-TAL effectors are presented). A core genome-based tree of 28 *X. translucens* strains is shown to the left side of the panel. Putative Type III secretion system effectors were identified via local Blastx against a database of known *Xanthomonas* effectors. Clustering of strains according to a hierarchical clustering of their encoded repertoires (Type III effector cluster dendrogram) is shown to the right side of the panel. Trees were visualized via FigTree, and mid-point rooted. Branches with bootstrap values below 98% are indicated in each tree. Xtt - *X. translucens* pv. translucens; Xtu - *X. translucens* pv. undulosa.

Bacterial organisms regularly secrete proteins and other molecules into the extracellular environment or directly into host cells to interact with their niche aiming at promoting growth, survival, colonization and virulence^14^. Many Gram-negative animal and plant bacterial pathogens, including *Xanthomonas spp.*, depend on a Type III secretion system (T3SS) for injecting effector proteins into the host cells to aid in pathogenesis^15,16^. Our previous work revealed that frequent loss and gain of a single gene encoding a Type II secreted cellulase effector, CbsA, promoted lifestyle switch for changes in tissue-specific host adaptation^2^. Here, we used a similar framework to define the evolution of host range shift in specialist and generalist subgroups (e.g. respectively, narrow and broad host range) of *Xanthomonas translucens*, arguably the most important re-emergent bacterial pathogen of cereals beyond rice^17–19^. We have determined that evolutionary loss and gain of Type III effectors (T3Es) govern the phenotypic switch between specialist and generalist behaviors of *X. translucens* subgroups for host range. Specifically, we determined that the broad host range lineage lost a single effector, XopAL1, to evade host defenses but at fitness cost on a primary host. Moreover, we assessed the potential for contemporary *X. translucens* specialist lineages to emerge via host jump, thus providing a framework for epidemiological tracking of particular lineages of risk.

## Results

### *Xanthomonas translucens* subgroups with differential host range behaviors encode significantly different T3E repertoires

Bacteria in the *X. translucens* species primarily infect grass and cereal plants in the Poaceae family, causing bacterial leaf streak (BLS) and blight, and black chaff of cereals^17,19^. Early documentation from the 1900s described the broad host range of generalist *X. translucens* pv. undulosa (Xtu)^20,21^. This subgroup uses a wide range of metabolites upon infection of host plants^22^ and causes disease in wheat, barley, rye, triticale, spelt, wild oat, ornamental asparagus, wild rice, and in a variety of wild grasses, totaling 12 plant genera (Fig. 1A and Fig. S1)^19–21,23–27^. In contrast, the closely related *X. translucens* pv. translucens (Xtt) is a specialist with limited metabolism that only causes disease in plants of the *Hordeum* genus (barley; Fig. 1A-B and Fig. S1)^20,22,24^. The main small grains affected by Xtt and Xtu are barley and wheat, respectively^17,24^. For example, upon spray inoculation for naturalistic entry and infection, Xtt and Xtu both caused water soaking symptom development in barley, but only generalist Xtu caused disease in wheat (Fig. 1B).

To better understand differences in niche specificity between the closely related (>97% average nucleotide identity)^28^ but phylogenomically distinct Xtt and Xtu (Fig. 1A,C), comparative genomic analyses were conducted and revealed that over 3,000 core proteins are shared between them (Table S1). On the other hand, each subgroup had over 100 unique conserved proteins (Table S2). Secreted proteins, which are often associated with virulence^14^, represented 3% and 3.5% of shared (core genome) and unique proteins (accessory genome), respectively (Tables S1-2). The number of unique, overall secreted proteins were not significantly different between Xtt and Xtu (Student’s *t*-test; *P* value = 0.7772). However, specialist Xtt has a larger T3E repertoire than generalist Xtu (Wilcoxon rank-sum test; *P* value = 3.436 e-06).

Based on the biological importance of T3Es^15,16^, we aimed to determine if there is a link between host range of *X. translucens* and encoded T3E repertoires. Hierarchical clustering of strains according to their encoded effector repertoires was congruent with the subspecies tree and with generalist and specialist behaviors (Fig. 1C). Specialist Xtt genomes encode an expanded effector repertoire that include XopAJ, XopAL1, XopE5, and HopW1 and an effector family expansion for XopP1 and XopP2. Only one unique T3E was identified in Xtu (XopE4), while seven were overall absent in this generalist subgroup (Fig. 1C). Moreover, Type III-secreted transcription activator-like (TAL) effectors were significantly more abundant in broad host range Xtu (Fig. S2; Student’s *t*-test; *P* value = 0.02511). Highly virulent Xtu strains have previously been found to precisely manipulate plant hormone pathways via the Tal8 (synonym TalDC) effector to cause wheat bacterial leaf streak^29^. This effector is not encoded by low-virulence strains of Xtu or any Xtt strain (Fig. S2)^29^. We therefore hypothesize the TAL effector family expansion correlates with adaptation to new niches as these are highly evolvable repetitive elements^30^. While we only focused on non-TAL effectors in this study, future work will explore host adaptation based on this special and abundant class of effectors.

### The T3E XopAL1 affects both host range and fitness in *X. translucens*

We hypothesized that individual T3Es might be responsible for host range differences between Xtt and Xtu. We first tested pathogenicity on both barley and wheat for Δ*hrcT* mutants that lack a critical component for the *X. translucens* T3SS by inoculating leaves via syringe infiltration^16^. Δ*hrcT* mutants of Xtt and Xtu did not cause symptoms in barley or wheat (Fig. S3A). Unlike the lack of symptom development upon natural infection via spray (Fig. 1B), Xtt caused chlorosis in wheat when inoculated via syringe infiltration (Fig. S3A). This has been described as a non-host or non-pathogenic response unlike the well-described hypersensitive response and is an undefined plant response that prevents disease development by Xtt^8,16,31–33^.

We then aimed to define if a unique Xtt effector prevented wheat pathogenesis by screening for loss of chlorosis. Individual deletions of the four Xtt-unique or single Xtu-unique effector genes did not alter pathogenicity in barley or chlorosis development in wheat upon infiltration inoculation except for *xopAL1* (Fig. S3). Notably, *xopAL1* deletion in Xtt produced water soaking symptom development in wheat similar to Xtu BLS (Fig. S3A). Moreover, complementation of the Xtt Δ*xopAL1* mutant strain or *xopAL1* introduction into the true wheat pathogen Xtu (Xtu miniTn7::*xopAL1*) triggered development of the non-pathogenic, chlorosis phenotype in wheat (Fig. S3). Introduction of the other three Xtt-unique effectors in Xtu did not trigger the wheat chlorotic response (Fig. S3B).

These findings were further assessed by spray inoculation, an infection method more similar to natural infections than syringe infiltration. Xtt Δ*xopAL1* caused BLS similar to Xtu on wheat and barley (Fig. 2A). On the other hand, introduction of this effector into Xtu disrupted its ability to cause disease in wheat (Fig. 2A). Curiously, deletion of *xopAL1* significantly decreased the symptomatic leaf area of barley inoculated via spray (Fig. 2B). Because of the reduced virulence of Xtt Δ*xopAL1* and Xtu although pathogenic is infrequently isolated from barley^34^, we suspected that XopAL1 plays a role in bacterial fitness beyond restricting the ability of Xtt to cause disease in wheat. Significantly higher endophytic populations of Xtt wild-type (WT) and Xtu miniTn7::*xopAL1* were observed in barley leaves in comparison to non-*xopAL1*-encoding strains (Xtt Δ*xopAL1* and Xtu WT; Fig. 2C). Moreover, Xtt Δ*xopAL1* had a similar population to Xtu in wheat upon syringe infiltration, whereas introduction of this effector in Xtu significantly reduced its population to similar levels as Xtt WT (Fig. S4). Altogether, these results demonstrated that *xopAL1* is a fitness factor for Xtt virulence and colonization in specialized host barley, but which restricts its ability to cause disease in wheat in an effector-dependent manner possibly via XopAL1 recognition.

**Fig. 2.**
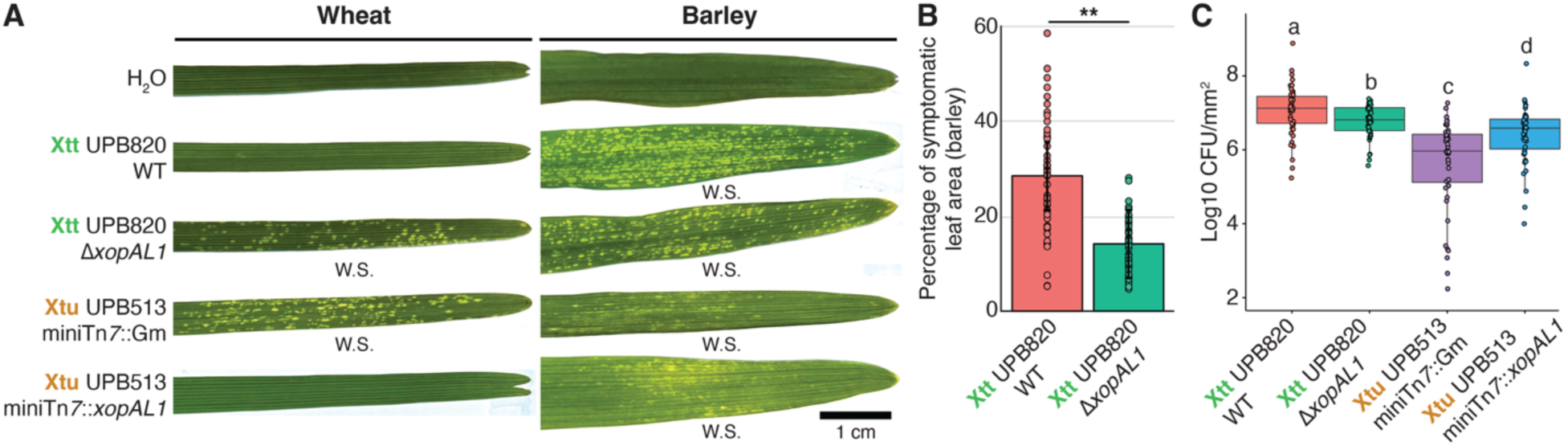
Deletion of the T3E gene *xopAL1* allows for host jump of Xtt from barley to wheat, while decreasing its pathogenic fitness in barley. **A.** Symptom development in barley and wheat leaves inoculated (spray) with Xtu and Xtt strains (wild-type, mutant, and insertion strains; OD_600_ 0.5) at six days post inoculation. Water soaking symptoms were visualized using a light box and appear as lighter areas in the leaf. Scale bar is shown in figure. **B.** Percentage of symptomatic leaf area of inoculated barley leaves (spray) with Xtt WT and Xtt Δ*xopAL1* shown in panel A. Original data of percentage of symptomatic leaf area (treatments; y-axis) was submitted to a transformation that combines the arcsine and square root functions. The Shapiro-Wilk test (W) was then applied to the transformed data, and it supported the assumption of normal distribution (W = 0.975; *P* value = 0.0671). ** indicates significant difference as determined by Student’s *t*-test (*P*<0.001; n = two independent replicates with 23 to 24 biological replicates each). **C.** Xtt and Xtu (wild-type, mutant, and insertion strains) populations in barley leaves inoculated via spray (OD_600_ 0.5) at 144 hours post inoculation. Different letters indicate significant difference as determined by Wilcoxon rank-sum test (*P*<0.05; n = two independent replicates with 23 to 24 biological replicates, each with three internal replicates). Wheat – *Triticum aestivum* L. genotype ‘Chinese Spring’; Barley – *Hordeum vulgare* L. cv. Morex; Xtt - *X. translucens* pv. translucens; Xtu - *X. translucens* pv. undulosa; W.S. – water soaking.

A cereal genotype diversity panel containing known susceptible hosts to Xtu was screened to assess if other plant species display a similar *xopAL1*-dependent resistance mechanism (Table S3). All screened hexaploid wheat genotypes (*Triticum aestivum* L., BBAADD genotype) presented a *xopAL1*-dependent resistance response against Xtt. Diploid (einkorn, AA genotype) and tetraploid (durum and emmer, BBAA genotypes) wheat genotypes were overall more susceptible to Xtt in a XopAL1-independent manner. Curiously, all tested Triticale lines (hexaploid genotype BBAARR) displayed a *xopAL1*-dependent resistance response against Xtt. Despite Triticale being a wheat-rye hybrid^35^, no *xopAL1*-dependent resistance was observed in rye, which remained resistant to Xtt. Curiously, genomic introduction of *xopAL1* into Xtu did not convert this subgroup into a non-pathogen of Triticale, as opposed to wheat. We therefore hypothesize that wheat resistance against Xtt may be mostly encoded by the DD sub-genome, which has the annual grass *Aegilops tauschii* Coss. as the donor species^36^.

A subset of this cereal genotype diversity panel was screened with individual mutant strains for the other Xtt-unique effectors (*xopAJ*, *xopE5*, and *hopW1*; Table S4). Our analysis revealed a *xopAJ*-dependent resistance response of oat (*Avena sativa* L., AACCDD genotype) against Xtt (Table S4; Fig. S5). Results thus suggest that this Xtt-unique effector determines its host range for causing disease in oat. Nevertheless, the downstream analyses described here focused mainly on XopAL1 due to its role in determining the ability of *X. translucens* to cause disease in its two main economic host plants, wheat and barley^17,24^.

### *xopAL1* has been lost in Xtu and presents an inherent risk for host jump by Xtt

Plant pathogen populations may shift and diversify their encoded virulence factor repertoires to avoid recognition by the host immune system^1,37^. We therefore tested if the emergence of broad host range Xtu correlated with the loss of *xopAL1*. Xtu was suggested to have diverged from a Xtt ancestor^38^, and we examined *xopAL1* evolution with maximum likelihood phylogenetic comparisons of whole genomes and the *xopAL1* locus neighborhood (Fig. 3A-B; Figs. S6-7), and by reconciling with the *xopAL1* tree (Fig. S8A). Specific neighborhoods of *xopAL1* were defined across the *X. translucens* species (Fig. 3A and Fig. S9). Our phylogenetic hypothesis testing of the *X. translucens xopAL1* locus determined that this effector was lost in Xtu (neighborhood type 1) during its divergence from a common ancestor with Xtt (neighborhood type 2; Fig. 3B-C). Loss of *xopAL1* was common across other *X. translucens* lineages. Three other inferred losses of *xopAL1* with acquisition of different genes replacing this effector were observed in subgroups with neighborhood type 4 (Fig. 3B). The origin of *xopAL1* (neighborhood type 2) was parsimoniously inferred in *X. translucens* to occur after divergence from a common ancestor with pathovar cerealis and strain ICMP 16317 in pathovar pistaciae (Xtp; neighborhood type 5; Fig. 3B). In addition, there was truncation of *xopAL1* in pathovar graminis (Xtg) by direct transposon insertion 5’ of the remaining sequence fragment (neighborhood type 3; Fig. 3B and Fig. S8B). For comparison, we analyzed the evolution of *xopE5* and *hopW1* (other Xtt-unique effector genes) and also observed loss in Xtu for both (Figs. S10-11, respectively).

**Fig. 3.**
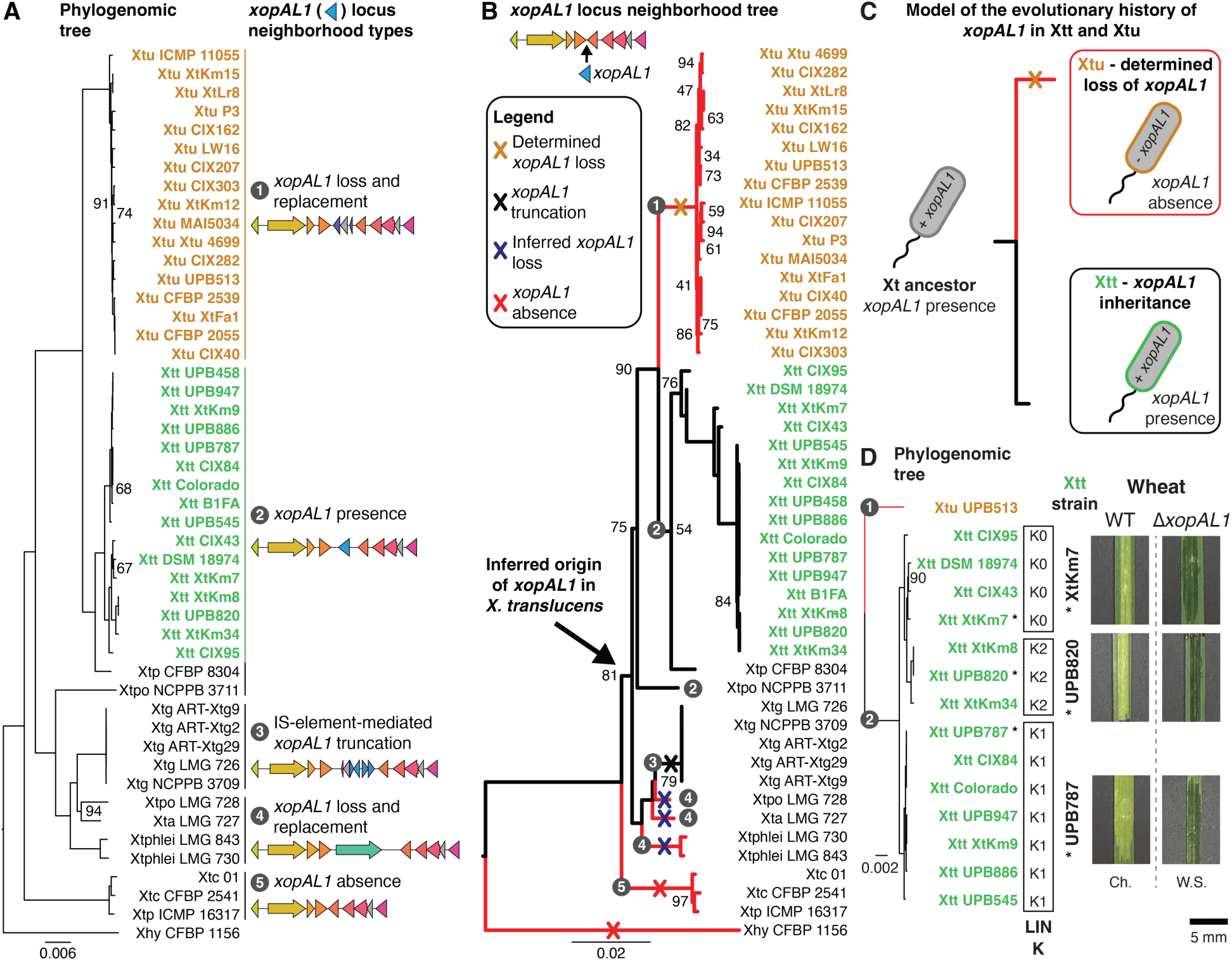
Loss of *xopAL1* promoted host jump of *X. translucens* pv. undulosa to wheat. **A.** Core genome-based tree of *X. translucens* strains and their *xopAL1* locus neighborhood types. A complete list of genes composing each neighborhood is shown in Fig. S9. **B.** Tree of the concatenated sequences of the genes flanking *xopAL1* in *X. translucens*. The *xopAL1* locus neighborhood types are indicated in each branch. Red-colored branches indicate subgroups that do not encode *xopAL1*, with determined and inferred loss or absence being shown. *X. hyacinthi* strain CFBP 1156 was included as an outgroup to root the trees in panels A and B. **C.** Model of the evolutionary history of *xopAL1* in Xtt and Xtu. **D.** Deletion of *xopAL1* promotes host jump of all Xtt genomic subgroups. A core genome-based tree including the K0, K1 and K2 genomic subgroups of Xtt is shown. Symptom development in wheat (*Triticum aestivum* L. genotype ‘Chinese Spring’) leaves inoculated (syringe infiltration) with representative Xtt strains (OD_600_ = 0.1) belonging to the three K genomic subgroups and their respective *xopAL1* mutants at three days post inoculation is shown to the right side of the panel. Scale bar is shown in figure. Trees were visualized via Figtree. Branches with bootstrap values below 98% are indicated in each tree. Ch. – chlorosis; W.S. – water soaking. *X. translucens* pathovars: Xtc – pv. cerealis; Xtp – pv. pistaciae; Xtg – pv. graminis; Xtpo – pv. poae; Xta – pv. arrhenateri; Xtphlei – pv. phleipratensis; Xtu – pv. undulosa; Xtt – pv. translucens; Xhy – *X. hyacinthi*.

Risk assessment is key for epidemic preparedness, but there is often a lack of precise knowledge about what genetic events lead to emergence, and about the potential of organisms to change their niche. Because loss of *xopAL1* allowed Xtu to jump host to wheat, we examined if deletion of this effector gene could lead to host jump for the three described, globally-distributed Xtt genomic subgroups (K0, K1, and K2)^34,39^. The deletion of *xopAL1* in representative K0, K1 and K2 strains led to host expansion to wheat (Fig. 3D). Xtt lineages are globally important in barley production systems^39^, and all have an inherent pathogenic capacity in wheat production systems based on *xopAL1* loss.

### *xopAJ* has been gained by Xtt and enhances virulence in barley

Gene gain is another mechanism for niche shift in *Xanthomonas*^2^. Using the same framework as for *xopAL1*, we determined that the oat-limiting effector gene *xopAJ* was originated in Xtt (neighborhood 2) after divergence with Xtu (neighborhood 1; Figs. S12-13). This gene was also gained in related pistachio-infecting *X. translucens* (Xtp) strain CFBP 8304 but further truncated via transposon insertion (neighborhood 3; Figs. S12-13). *xopAJ* is not encoded by any other *X. translucens* subgroup and is not encoded by Xtt in the same locus as other *xopAJ*-encoding *Xanthomonas* spp. (Fig. S14). Moreover, Xtt-encoded *xopAJ* is more phylogenetically closely related to *xopAJ* encoded by a distant species, *Paracidovorax citrulli*, than to *xopAJ* encoded by other *Xanthomonas* spp. (Fig. S15). These results suggest that *xopAJ* evolution is dynamic and was gained in Xtt by horizontal gene transfer but lost by dicot pathogen Xtp. We hypothesized that gain of *xopAJ* by Xtt increases its adaptive fitness to colonize barley. Deletion of *xopAJ* in Xtt significantly decreased the symptomatic leaf area of barley upon spray inoculation (Fig. S16A-B), but no significant changes were observed for endophytic populations of the pathogen (Fig. S16C). Altogether, results show that gain of *xopAJ* enhanced virulence of Xtt in barley, while not affecting fitness for plant colonization.

### XopAL1 mediates global transcriptional reprogramming in wheat leading to a chlorotic non-host response

The basis of chlorosis development in a non-host response is widespread in *Xanthomonas*-plant interactions but remains unclear^8,31,32^. For instance, a distant XopAL1 ortholog encoded by *Xanthomonas campestris* pv. campestris (Fig. S8) also triggered chlorosis development in wheat when transformed in Xtu (Fig. S17). We thus carried out RNA sequencing (RNA-seq) to elucidate the mechanistic basis of non-host response that restricts bacterial infection. Expression profiles were compared between wheat leaves inoculated with *xopAL1*-encoding strains (Xtt WT and Xtu miniTn7::*xopAL1*), non-encoding strains (Xtt Δ*xopAL1* and Xtu WT), and mock-inoculated leaves. A principal component analysis (PCA) of the RNA-seq count data demonstrated that samples clustered in two primary transcriptional profiles in a *xopAL1*-dependent manner, reflecting the ability of strains to cause disease in wheat or not (Fig. 4A). Accordingly, much higher numbers of significantly differentially expressed genes (DEGs) were identified in plants inoculated with Xtt and Xtu encoding *xopAL1* when compared to mock-treated samples (12,397 and 9,118 DEGs, respectively; Table S5) and in comparison to non-encoding strains of these subgroups (607 and 285 DEGS, respectively; Table S5) (Fig. 4B).

**Fig. 4.**
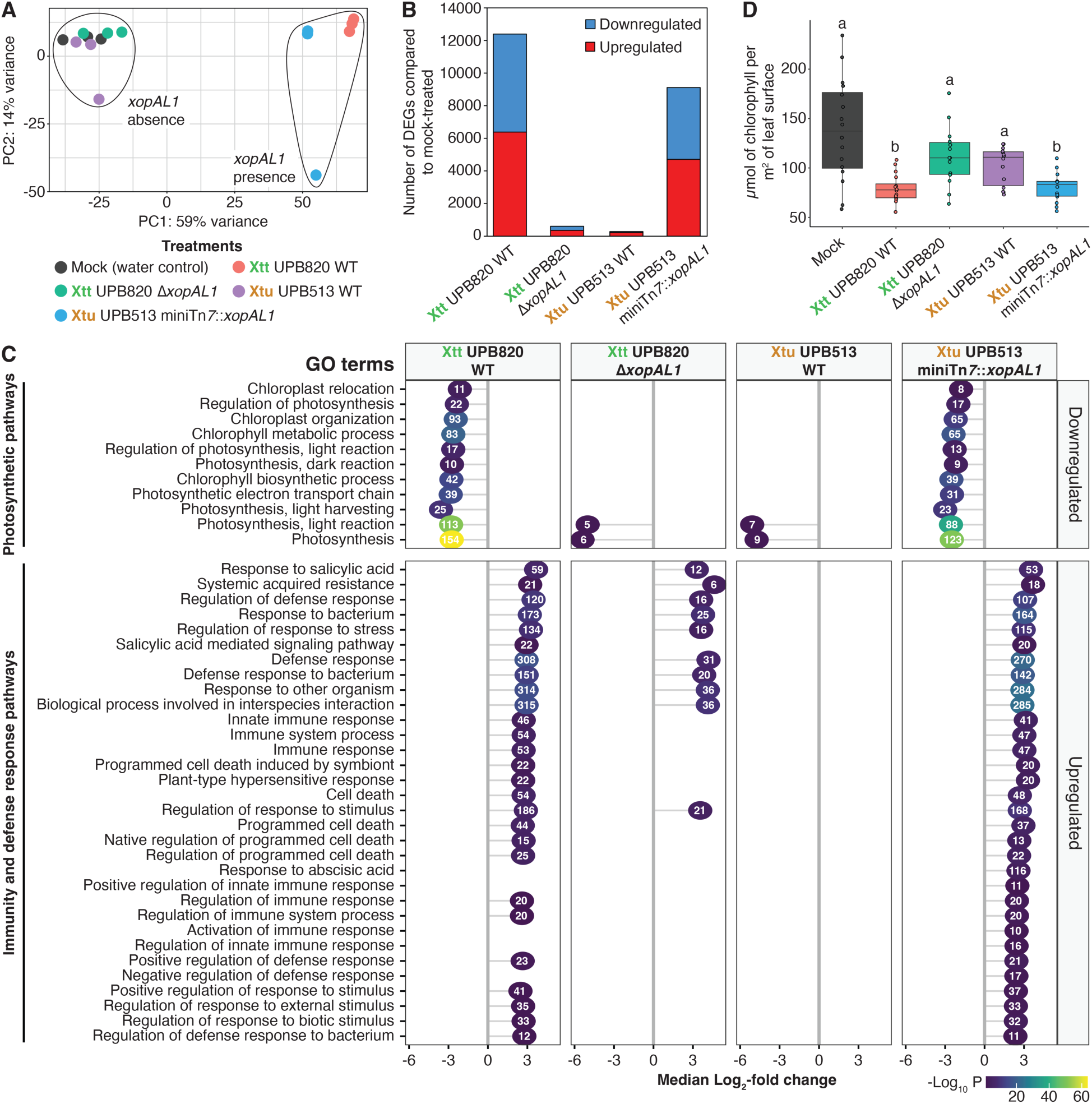
XopAL1 mediates global transcriptional changes in inoculated wheat plants, leading to a chlorotic non-host response. **A.** Principal component analysis (PCA) of the normalized RNAseq data transcripts per million of wheat leaf tissue infected with different *X. translucens* strains and mock (treatments). Ellipses are arbitrarily drawn to indicate clustering of treatment based on *xopAL1* presence and absence. **B.** Number of significantly differentially expressed genes (DEGs; downregulated and upregulated) in wheat from each bacterial treatment compared to mock. **C.** A subset of Gene Ontology (GO) terms significantly enriched in wheat from bacterial treatments versus mock comparisons. The number of genes that fall into a GO term from each comparison are indicated within each dot. *P* values are indicated by dot color (color scale is shown in figure). Upregulated and downregulated GO terms are indicated in the plot. **D.** Chlorophyll concentration measurement in wheat leaves (*Triticum aestivum* L. genotype ‘Chinese Spring’) inoculated with Xtt and Xtu strains (wild-type, mutant, and insertion strains) via leaf syringe infiltration (OD_600_ 0.1) at three days post inoculation. Different letters indicate significant difference as determined by Wilcoxon rank-sum test (*P*<0.05; n = two independent replicates with 6 to 9 biological replicates, each with three internal replicates). Xtt - *X. translucens* pv. translucens; Xtu - *X. translucens* pv. undulosa.

A gene ontology (GO) term enrichment analysis was performed to investigate the specific plant processes in wheat that led to the non-host response. Our analysis revealed that upregulated DEGs in response to inoculation with Xtt and Xtu expressing *xopAL1* were significantly enriched in GO terms associated with immunity, induced defense response, defense signaling, and programmed cell death (Fig. 4C; Table S5). On the other hand, significant overrepresentation of GO terms associated with photosynthetic pathways, including chloroplast and chlorophyll processes, were observed among the downregulated DEGs in response to strains expressing *xopAL1* (Fig. 4C; Table S5). Based on this, we hypothesized that chlorosis is the result of reduced photosynthetic activity and chlorophyll content in wheat upon XopAL1 recognition. Indeed, we observed significantly lower chlorophyll concentration in wheat leaves inoculated with *xopAL1*-encoding Xtt and Xtu in comparison to those inoculated with non-encoding strains and mock-inoculated leaves (Fig. 4D). These findings indicate that recognition of XopAL1 in wheat results in extensive transcriptional changes that ultimately lead to a non-host type of resistance response.

## Discussion

Emerging infectious diseases often result from microbial niche shift. Multiple genetic mechanisms control emergence including gene loss and gain. Genome and episome reduction are strongly associated with host specialization and gain can promote adaptation to a particular environment or resist control interventions (e.g. antibiotics)^40,41^. Here, were have determined that loss and gain of single T3Es control specialist and generalist behaviors of Xtt and Xtu for host range. Specifically, our results collectively demonstrate that a) XopAL1 is both a fitness and virulence requirement for Xtt; b) this effector is a determinant of host range since its loss contributed for host expansion of Xtu to wheat by circumventing an otherwise effector-triggered defense response of this host plant; and c) gain of XopAJ by Xtt limited its ability to cause disease in oat but contributes for full virulence in barley.

Bacterial pathogens commonly colonize host plants by delivering secreted factors that alter plant cellular processes to promote virulence, but these same factors can also trigger plant immune responses that restrict infection if recognized by cognate plant resistance (R) proteins^15,16^.

Absence or presence of specific T3Es dictates host range of plant pathogens in an effector-triggered immunity (ETI) -dependent manner. Examples of effectors affecting host range include the XopAG, XopQ, and AvrBsT effectors of *Xanthomonas* spp.^9,10,12,32^, and HopQ1-1 effector of *Pseudomonas syringae*^11^. Moreover, effectors have demonstrated fitness requirements for leaf colonization in particular hosts^37,42^. For instance, absence of AvrBsT expands the host range of *Xanthomonas perforans* while decreasing the fitness of this pathogen in tomato fields^42^.

These studies demonstrate how T3E repertoires may vary in response to resistance within individual cropping or model plant systems but lack an evolutionary framework. Here, we determined the phylogenetically-supported evolutionary history and ecological function of T3E loss and gain events related to fitness and host range. Using the generalist model Xtu, we posit that *xopAL1* was one of multiple loss events that led to emergence of this pathogenic generalist subgroup. Xtu has lost other niche-specific factors for tissue-specificity. Our previous work demonstrated that loss of the vascular-associated pathogen gene *cbsA* is frequently associated with non-vascular tissue colonization. This gene is encoded by the vascular, specialist Xtt but was lost in non-vascular, generalist Xtu via transposon insertion^2^. Here, we have also observed truncation of *xopAL1* in Xtg, and *xopAJ* in Xtp via transposon insertion. Transposons shape microbial niche-specificity, and these insertion sequence (IS) elements are major drivers of evolutionary ecological shifts of bacterial and fungal species^2,43–45^. Overall, we hypothesize that gene loss, frequently mediated by IS elements, is a common strategy for niche shifts across behaviors.

Interestingly, XopAJ, which we have determined here as a host range-limiting factor for *X. translucens* in oat, was gained in Xtt and contributes for virulence in barley. We speculate that gain of effectors upon host specialization provides resource access on a preferred host. XopAJ (synonym of AvrRxo1) was initially described as a host range-limiting factor of the rice pathogen *X. oryzae* pv. oryzicola (Xoc) when inoculated into maize lines carrying the cognate resistance gene Rxo1^46^. This effector has been shown to suppress pattern-triggered immunity (PTI) responses of host and non-host plants^47^, and to enhance virulence of Xoc^48^. Xtt is limited to barley (*Hordeum* spp.), and it remains to be determined whether XopAJ displays similar molecular functional roles for host colonization and pathogen virulence as observed in other *Xanthomonas* spp.

Hosts respond dramatically different to specialized and generalist pathogens and could be used to inform resistance trait discovery. Components of the wheat defense response pathway identified here via RNA-seq compose a valuable pool of candidates to identify the interactor(s) of XopAL1 and generate resistance in other hosts including barley. Moreover, we observed downregulation of chloroplast processes, which play a central role in plant immunity^49^. Together, our RNA-seq results help to understand the basis of the chlorotic non-host response that includes upregulation of defense responses with downregulation of photosynthetic processes.

Preparedness for mitigation of new epidemics, especially when host jump occurs, depends on strong knowledge of the evolution of niche shift. The research described here provides a framework for reconstructing the evolutionary history of host jump in an important cereal pathogen and allows us to predict pathogen emergence based on gene loss and gain. Based on our results, we developed a tested evolutionary model for *X. translucens* intergenera host jump centered on effector gene loss and gain (Fig. 5). Xtu is a generalist that has fewer T3Es, and loss of *xopAL1* promoted its broad host range behavior. Conversely, gain of *xopAJ* in the specialist Xtt restricted its ability to cause disease in oat. A practical implication of our model is that management of BLS of wheat and oat must include surveillance of all lineages of Xtt in addition to Xtu due to their inherent risk for jumping hosts based on loss of single genes. Ultimately, this work demonstrated that the shift between specialist and generalist subgroups of *X. translucens* for host range is modulated by presence and absence of single genes, with gene loss and gain being major drivers for changing niches and ecological behaviors of pathogens.

**Fig. 5.**
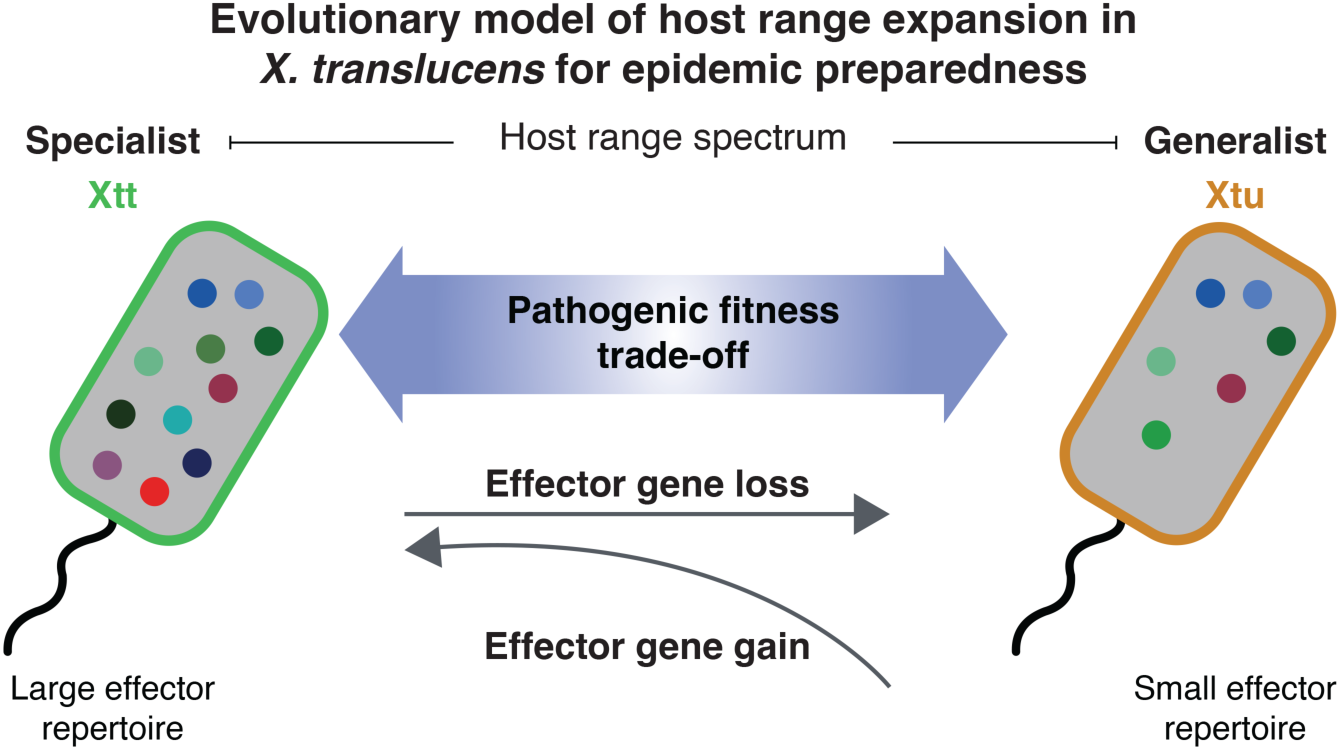
Proposed evolutionary model of host range expansion in *Xanthomonas translucens* based on effector gene loss and gain. Taken together, phenotypic and phylogenetic analyses support a model where loss of a single Type III secretion system effector gene, *xopAL1*, allowed for host range expansion to wheat by the generalist Xtu, while gain of *xopAJ* in the specialist Xtt restricted even more its host range breadth. Our model also implies that Xtt strains have an inherent risk for host jump via individual losses of *xopAL1* and *xopAJ*, which has profound effects for epidemic preparedness and management practices. Circles with different colors in figure represent different effector repertoires encoded by Xtt and Xtu. Xtt - *X. translucens* pv. translucens; Xtu - *X. translucens* pv. undulosa.

## Materials and Methods

### Bacterial strains, plasmids, and culture conditions

Bacterial strains and plasmids used in this study are listed in Table S6. *Xanthomonas* strains were recovered from -80°C glycerol stocks and grown for two days at 28°C on nutrient agar (NA) or nutrient broth (NB) (3 g/L beef extract, 5 g/L peptone, 15 g/L agar). When needed, the antibiotics kanamycin (Km), gentamicin (Gm), or cycloheximide (CHX) were used at concentrations of 50 µg/ml, 5 µg/ml, and 50 µg/ml, respectively. Sterile deionized water was used for suspending cells in liquid. *Escherichia coli* strains were cultured in Luria-Bertani (LB) medium at 37°C. LB was supplemented with kanamycin (50 µg/ml), gentamicin (10 µg/ml), or ampicillin (100 µg/ml) whenever necessary. *E. coli* WM3064, a diaminopimelic acid (DAP) auxotrophic strain used for conjugation transformation, was grown as described above, but with LB medium supplemented with DAP (200 µM).

### Recombinant DNA techniques

Genomic DNA of respective strains was isolated using the Monarch Genomic DNA Purification kit (New England Biolabs). Polymerase chain reactions (PCR) were performed using Q5 High-Fidelity DNA Polymerase (New England Biolabs) for cloning sequences into plasmid vectors of interest or using Quick-Load *Taq* 2X Master Mix (New England Biolabs) for other processes. PCR reactions were carried out using standard manufacturer protocols in a ProFlex PCR System thermal cycler (Applied Biosystems). PCR products and agarose gel fragments were purified using the QIAquick PCR and Gel Cleanup kit (QIAGEN). Competent cells of *E. coli* strains were made and transformed using the Mix & Go! *E. coli* Transformation Buffer set (Zymo Research). Plasmid DNA was isolated from overnight cultures of *E. coli* using the QIAprep Spin Miniprep kit (QIAGEN).

Transformation of Xtt UPB820 and Xtu UPB513 with respective plasmids (Table S6) was performed by electroporation (1.8 kV, 200 Ω, 25 μF and a time constant lower than 5 ms). Some Xtt UPB820 strains were transformed via electroporation with pPneo-GFP for kanamycin resistance and GFP expression (Table S6). Transformation of Xtt UPB787 and XtKm7 was made via conjugation using the *E. coli* donor strain WM3064. In summary, Xtt strains were grown overnight in 40 mL of NB (28 °C, 220 rpm), and *E. coli* WM3064 strains were grown overnight in 20 mL of LB + DAP + respective antibiotic (37 °C, 220 rpm). After, Xtt and *E. coli* cells were collected by centrifugation (4,000 rpm for 10 minutes), suspended into the same volume of media (40 mL of NB for Xtt and 20 mL of LB + DAP for *E. coli* WM3064), centrifuged again and suspended with adequate volume of media to obtain an optical density at 600nm (OD_600_) of 3.0. Then, Xtt and *E. coli* cells were combined at a 1:1 ratio and 200 µL aliquots were spotted on top of MF-Millipore MCE membranes (25 mm diameter, 0.45 µm pore size; Merck Millipore Ltd.) overlaid on NA + DAP. Eight membranes were spotted per transformation event. Plates were incubated for 24 hours at 28°C, and membranes were then recovered by using sterile forceps and placed into conical tubes containing 20 mL of NB. Cells were removed from membranes by vortexing on maximum for 1 minute, and membranes were removed using sterile forceps. At last, cells were collected by centrifugation (4,000 rpm for 10 minutes), suspended in 1 mL of NB, diluted by 10-fold serial dilution until 10^-2^ and 500 µL aliquots of the 10^-1^ and 10^-2^ suspensions were then plated on NA + antibiotic selective medium (150 mm diameter plates) for growth of transformed cells.

Clean deletion of genes of interest (GOI) in strains of *X. translucens* (Table S7) was made via *sacB* counter-selection using the pK18mobsacB plasmid, as previously described^2^. Clean deletion of each GOI was confirmed via PCR (Table S8). Complementation of *xopAL1* and *xopAJ* in the respective mutant strains of each in Xtt UPB820 was also performed via the pK18mobsacB plasmid. The upstream and downstream regions of each gene, together with their open reading frames, were amplified altogether using the upstream region forward primer and the downstream region reverse primer (Table S8) and then cloned into pK18mobsacB at the HindIII restriction site via Gibson Assembly (New England Biolabs). Insertion of *xopAL1* and *xopAJ* back into their original loci was confirmed via PCR (Table S8).

To obtain the constructs for genomic insertion via miniTn*7*, the open reading frames of each GOI containing promoter and terminator regions were amplified by PCR and then fused (when needed) and cloned into pUC18miniTn*7*T-Gm at the HindIII restriction site via Gibson Assembly (New England Biolabs). The obtained constructs were confirmed via PCR (Table S8) and transformed into Xtu UPB513 together with pTNS3 (Table S6) to promote transposition and single gene insertion. Transformant cells were selected by plating on NA + Gm and confirmed for GOI genomic insertion by PCR (Table S8).

The *xopAL1*_Xcc8004_ gene (400-bp promoter plus 100-bp terminator, amplicons LM169-170 and LM171-LM172; Table S8) was cloned from *X. campestris* pv. campestris (Xcc) strain 8004 by GoldenGate strategy using *Bsa*I-mediated restriction-ligation into pEL667, a GoldenGate-compatible derivative of pBBR1MCS2. Ligation products were introduced into *E. coli* by electroporation and conjugated into Xcc using pRK2073 as a helper plasmid in triparental matings.

### Plant growth conditions and inoculation methods

Barley (*Hordeum vulgare* L. cv. Morex) and wheat (*Triticum aestivum* L. varieties ‘Chinese Spring’, RB07, Glenn, and AGS2055) seeds were germinated and grown in growth chambers with cycles of 16 hours of light per day at 22° to 24 °C and 70% relative humidity for leaf syringe infiltration inoculations, or in greenhouse conditions for temperature, relative humidity and light for spray inoculation. In both experiments, seeds were germinated directly in potting mix (PRO-MIX BX General Purpose; Premier Tech Growers and Consumers Inc.) in 3.5 inches diameter square pots (four seeds per pot; Kord Products).

For spray inoculation, the primary leaf (counting from the bottom) of seven-day-old wheat and barley plants was sprayed till run off on the abaxial and adaxial surfaces with water-based bacterial inocula (OD_600_ of 0.5) or sterile deionized water as mock control. After leaves were dry, pots with plants were transferred to trays filled with approximately 0.5 inch of tap water. Trays were then covered for four days to promote high relative humidity, which facilitates infection via spray inoculation. Plants were kept for two additional days in greenhouse conditions after removing the covers and development of water soaking symptoms was evaluated at six days post inoculation. Individual inoculated leaves were taped to transparency film sheets and pictures were taken using a LED light box (Tikteck). When appropriate, the total and symptomatic leaf areas were measured via the ImageJ software^50^ by using a set scale. A standardized length of 5 cm from the top to bottom of each inoculated leaf was used for individual measurements. The percentage of symptomatic leaf area was then calculated using the obtained areas. The experiment was conducted independently twice, with 23 to 24 independent plants being inoculated for each treatment in each independent replicate.

For bacterial population quantification in barley following spray inoculation, leaves inoculated as described above were collected at six days post inoculation. The leaves were surface-disinfested by soaking into 70% ethanol for five seconds, followed by two consecutive rinses in sterile deionized water for five seconds each. This was done so that the isolation targeted those bacterial cells that naturally moved and multiplied into the inner leaf tissue. After, two leaf discs were collected at each side of the main leaf vein using a standardized length of 2.5 cm from the top to bottom of each inoculated leaf by using a 2.5 mm diameter punch (Acuderm Inc.). Leaf discs were then flash frozen in liquid nitrogen and stored at -80°C until processed. For bacterial isolation, leaf discs were ground in 100 µL of sterile deionized water in 2-mL microcentrifuge tubes containing a sterile 5 mm diameter stainless steel bead (QIAGEN) using the TissueLyzer II apparatus (30 Hz for 1 minute; QIAGEN). Samples were then transferred to 96-well plates (Thermos Scientific), diluted by 10-fold serial dilutions and plated as 5 µL drops in triplicates in selective media (NA + CHX + Km for Xtt strains transformed with pPneo-GFP, and NA + CHX + Gm for Xtu strains with miniTn*7* genomic insertion). Colony forming units (CFU) were enumerated after three days of incubation at 28 °C. The experiment was conducted independently twice, with 23 to 24 independent plants being inoculated for each treatment in each independent replicate.

For infiltration inoculation, the first two leaves (counting from the bottom) of fourteen-day-old plants (wheat and barley) were inoculated using needleless syringes. Bacterial strains were suspended in sterile deionized water (OD_600_ of 0.1) for inoculation. Control plants (mock) were inoculated with sterile deionized water. Three to four independent plants were inoculated for each treatment, and experiments were conducted independently at least twice. Development of symptoms was recorded three days post inoculation (dpi), in which lesions associated with pathogenicity were recorded (water soaking – disease development; chlorosis – non-host response; no symptom – no pathogenicity). Pictures of inoculated leaves were taken at 3 dpi by placing them on top of a black non-reflective photography display board (Neewer) and using the camera of a cell phone mounted on a tripod (UBeesize) with a LED cell phone circle light (QIAYA). Germination, growth, inoculation, and evaluation of the plant species used to screen a cereal genotype diversity panel (Tables S3-4) with Xtt and Xtu strains were performed as described for infiltration inoculation.

Inoculations for bacterial growth curves performed in planta were made via leaf syringe infiltration, as described above, but using a lower optical density of each bacterial suspension (OD_600_ of 0.001). Plants were inoculated in triplicates per time point and treatment, and samples were collected at 2, 24, 72, 120, and 168 hours post inoculation using a 2.5 mm diameter punch (Acuderm Inc.), flash frozen in liquid nitrogen and stored at -80 °C until processed. Bacterial isolation, serial dilution, plating, and enumeration were performed as described above. Experiments were performed independently twice.

### Whole genome sequencing

Genomic DNA for Xtt strains Colorado, CIX84, UPB545, UPB787, and UPB820, and Xtu strain UPB513, was extracted using Genomic-tip 20/G and the Genomic DNA Buffer Set (QIAGEN) and then sequenced as previously described^51^. Briefly, PacBio long-read, single-molecule real-time sequencing was used to sequence extracted genomic DNAs, and assembly was performed via Flye v2.4^52^ for strains CIX84 and UPB545, and via HGAP v4 (PacBio) for strains Colorado, UPB787, UPB820, and UPB513 using default settings. Whole genomes were deposited at GenBank under accession numbers CP159781 for Xtt UPB545, CP159782 for Xtt CIX84, and JBEFAI000000000 for Xtu UPB513 (Table S9). Availability of the genomes of Xtt strains Colorado, UPB787, and UPB820 is shown in Table S9.

### Pangenome analyses to identify core and unique proteins among Xtt and Xtu

Pangenome analysis to identify core and unique proteins encoded by all analyzed Xtt and Xtu strains (Tables S1-2 and Table S9) was performed by comparing the gene presence/absence output matrix obtained when analyzing genomes via Panaroo v1.3.3, a tool designed to perform core and pangenome comparisons of prokaryotes^53^. Core proteins are determined here as proteins encoded by all strains in both pathovars translucens and undulosa. On the other hand, unique proteins of a specific pathovar are determined here as proteins encoded by all strains of a specific pathovar but are not encoded by any strain of the other pathovar. The proteome of a representative strain of each pathovar was searched for secretion signals using SignalP v6.0 to predict secreted proteins^54^. Proteomes were obtained by annotating genomes via Prokka v1.14.6 (see below)^55^. SignalP results were then used to identify which core and unique proteins of Xtt and Xtu are predicted to be secreted. Proteins in which a secretion signal was identified but were predicted to locate to the outer membrane or periplasm were excluded when calculating the percentage of secreted proteins within core and unique proteins. Additional genes encoded uniquely by subsets of strains belonging to one pathovar or the other were also identified and analyzed as described above to compare percentages of uniquely secreted proteins in each pathovar via statistical analysis (described below). Hypothetical proteins were not considered to calculate the proportion of secreted proteins in the core genome but were considered when performing this analysis in genes found uniquely in pathovars translucens or undulosa. We manually curated secretion of proteins annotated as hypothetical by searching for encoded functions using the blastp algorithm (<https://blast.ncbi.nlm.nih.gov/Blast.cgi?PAGE=Proteins>) and retrieving the best hit for each searched protein.

### Phylogenomic and phylogenetic analyses

Phylogenomic trees depicting the evolutionary relationships of examined *Xanthomonas* species for host range comparisons (Fig. 1A and Fig. S1) were constructed from well-supported 1780 single-copy orthologous genes (SCOs) trees. SCOs were determined via OrthoFinder^56^, and individual SCO phylogenies were constructed by first aligning via *mafft* v7.487^57^, trimming via *ClipKIT* v1.3.0^58^, and building the phylogeny via *IQ-TREE* v2.2.6 with 1,000 ultrafast bootstrap iterations^59^. Individual SCO trees with at least 75% average node support were concatenated for a multigene phylogenomic tree by applying best predicted evolutionary models to each SCO partition.

All other whole-genome phylogenies were performed based on core genome alignment. Genomes of interest were annotated via Prokka v1.14.6 specifying the appropriate genus and species with the “--genus” and “--species” options, respectively, and otherwise with default settings^55^, followed by core genome alignment using Panaroo v1.3.3 with a core threshold of 0.98 and option “—clean-mode strict”^53^. Alignments were then filtered for recombination by first using maximum likelihood recombination inference through ClonalFrameML v1.12^60^ followed by alignment masking with maskrc-svg v0.5 (https://github.com/kwongj/maskrc-svg), and trees were built using IQ-TREE v2.2.2.7 with 5,000 ultrafast bootstrap replicates.

A phylogenetic tree for the *xopAL1* locus neighborhood, which includes four upstream and five downstream genes of *xopAL1*, was constructed by first running Parsnp v1.7.4^61^ to align the nucleotide sequence of this neighborhood from the reference strain Xtt UPB820 (extracted with the GetFasta command from the BEDTools suite)^62^ with the full genomes of the other selected assemblies (Table S9), then converting the Parsnp output to a FASTA alignment using Harvesttools v1.2^61^, and finally inferring a tree with IQ-TREE v2.2.2.7 using 5,000 ultrafast bootstrap replicates. The same method was used to generate phylogenetic trees of the *xopE5* (six upstream and five downstream genes) and *hopW1* (five upstream and seven downstream genes) loci neighborhoods. An amino acid sequence-based phylogenetic tree of the *xopAJ* locus neighborhood (six upstream and four downstream genes of *xopAJ*) was generated by identifying orthologs of each of these genes in the genomes of interest via OrthoFinder v2.5.5, aligning them individually with MAFFT v7.520 using default parameters. Poorly aligned regions were removed from alignments using Trimal v1.5.0 using the ‘-automated1’ method^63^. A concatenated locus tree was inferred using IQ-TREE v2.2.2.7 with 1,000 ultrafast bootstrap replicates and gene concordance factors were calculated for each node.

Phylogenetic trees for individual genes within the *xopAL1* locus neighborhood were constructed by first identifying orthologs to these genes in Xtt UPB820 via OrthoFinder v2.5.5, extracting the focal sequences with BEDTools GetFasta, aligning them with MAFFT v7.520, and inferring trees using IQ-TREE v2.2.2.7 with 1,000 ultrafast bootstrap replicates. *xopAL1* encoded by the outgroup representative *Ralstonia solanacearum* strain 10314 was included for the phylogenetic tree of *xopAL1*. The phylogenetic trees of *xopE5*, *hopW1*, and *xopAJ* were constructed via the same method but using *Paraburkholderia nemoris* strain RL18-011-BIC-A and *Ralstonia mannitolilytica* strain LMG 18090 as outgroups for *xopE5*, *Pseudomonas amygdali* strain CFBP 1650 and *Erwinia psidii* strain LPF 534 as outgroups for *hopW1*, and *Paracidovorax citrulli* strains AAC00-1 and DSM 17060 as outgroups for *xopAJ*.

Classification of strains of *X. translucens* pv. translucens in the K0, K1, and K2 sub-genomic groups, as determined by life identification numbers (LINs) calculated with LINbase, was performed as previously described^39^.

### Identification of Type III secretion system effector repertoires

The effector repertoires of strains of *X. translucens* were identified as previously described via BLAST 2.8.1+ blastx algorithm^39^, excluding transcription activator-like effectors (TALEs). Briefly, a database of known *Xanthomonas* spp. Type III secretion system effectors (https://euroxanth.ipn.pt/doku.php?id=bacteria:t3e:effectors) was used as subject to search the query genomes. Results were filtered for hits with coverage of over 200 amino acids and percent amino acid identity of at least 60%. TALEs were identified and classified using AnnoTALE v1.5^64^. The effector repertoire heat map and the Type III effector cluster dendrogram (hierarchical clustering) figures were generated using the ComplexHeatmap^65^ and RColorBrewer (https://cran.r-project.org/web/packages/RColorBrewer/index.html) packages in R v4.1.0^66^.

### RNA-sequencing and analysis

To determine wheat response to XopAL1, an RNA-seq analysis was performed. Wheat plants were grown as described above and inoculated with Xtt UPB820 WT, Xtu UPB513 WT, Xtt UPB820 Δ*xopAL1,* Xtu UPB513 miniTn*7*::*xopAL1,* or a water (mock) control. Each treatment had three independent replicates, and bacterial inoculations were performed at an OD_600_ of 0.001. Tissue was collected 24 hours post inoculation, flash frozen in liquid nitrogen, and homogenized using a sterile 5 mm diameter stainless steel bead (QIAGEN) using the TissueLyzer II apparatus (30 Hz for 1 minute; QIAGEN). Total RNA was extracted using TRIzol (Thermo Scientific) and DNA was removed using the TURBO DNA-free kit (Thermo Scientific) following manufacturer’s guidelines. Plant and 16S ribosomal depletion was performed and total RNA libraries were constructed using random oligos. RNA libraries were multiplexed and sequenced using a NextSeq 2000 (Illumina) with 150-bp paired-end reads and a target of 30 million reads per sample.

RNA-seq analysis was performed using the Nextflow/nf-core “rnaseq” pipeline v3.10.1^67^, using the *Triticum aestivum* L. cv. ‘Chinese Spring’ reference genome (version 2.1) obtained from Phytozome. RNA-seq data were processed using the nf-core RNA-seq pipeline implemented with Nextflow. This pipeline included adapter trimming with TrimGalore! v0.6.5 (<https://www.bioinformatics.babraham.ac.uk/projects/trim_galore/>), alignment of reads to the wheat genome with STAR v2.7.10b^68^, and transcript abundance estimation with Salmon v1.10.1^69^. Quality metrics for each step were aggregated using MultiQC v1.14^70^. The read count Table output by the pipeline was used to identify differentially expressed genes using the DESeq2 package^71^. Genes with an adjusted *P* value (FDR) < 0.05 were considered significantly differentially expressed. Genes with a log₂ fold change (log₂FC) > 0 were defined as upregulated, while those with log₂FC < 0 were classified as downregulated. The raw RNA-seq fastq files are available at GenBank under accession number GSE299780.

Normalized gene counts produced from DESeq2 were used to generate a principal component analysis (PCA) plot using ggplot2 in R^72^. Gene ontology (GO) analysis was performed to identify enriched biological functions among differentially expressed genes. GO term annotations for each gene in the wheat ‘Chinese Spring’ reference genome were obtained from Phytozome (<https://phytozome-next.jgi.doe.gov>). Enrichment analysis was conducted separately for upregulated and downregulated genes using the Gene Ontology Resource (<http://geneontology.org>)^73–75^. Only significantly overrepresented GO terms (adjusted *P* value < 0.05) were considered enriched among downregulated or upregulated genes. Selected GO terms related to plant defense and photosynthesis were visualized using a Cleveland dot plot generated in R with ggplot2.

### Chlorophyll concentration measurement in wheat leaves

Xtt and Xtu strains were suspended in sterile deionized water (OD_600_ of 0.1) and inoculated into fourteen-day-old wheat leaves (*Triticum aestivum* L. genotype ‘Chinese Spring’; second leaf counting from the bottom) using needless syringes as described above for infiltration inoculation. Control plants (mock) were inoculated with sterile deionized water. At three days post inoculation, leaves were collected and the chlorophyll concentration in inoculated areas was measured in triplicates in each leaf using the MC-100 Chlorophyll Concentration Meter (Apogee Instruments, Inc.). Chlorophyll concentration was measured as micromoles per square meter (µmol m^-2^) of leaf using the manufacturer’s specific settings for wheat and the standard operational protocol for the instrument. The experiment was conducted independently twice, with six to nine independent plants being inoculated for each treatment in each independent replicate.

### Identification and visualization of genomic neighborhoods

To identify the *xopAL1*, *xopAJ*, *xopE5*, and *hopW1* genomic neighborhoods in *Xanthomonas* spp., Prokka-annotated genomes were aligned and visualized via progressiveMauve using default settings^76^ to determine flanking genes and neighborhoods. Genomes were then uploaded to the Benchling cloud-based platform (https://www.benchling.com) to extract the sequences of genomic neighborhoods in GBK format, which were submitted to cluster comparison analysis via Clinker v0.0.23 (default settings) to generate figures^77^.

### Protein alignment and visualization

Amino acid sequence alignment of representative sequences of XopAL1 within *Xanthomonas* spp. was performed via the T-Coffee Multiple Sequence Alignment Server^78^ and visualized using pyBoxShade (https://github.com/mdbaron42/pyBoxshade). Percent coverage and similarity were determined via the blastp algorithm.

### Data analysis and visualization

Data from number of proteins encoded by Xtt and Xtu strains (total and unique proteins), and data from number of TAL effectors encoded by both of these subgroups were compared by two-tailed Student’s *t*-test. Data from number of proteins in the effector repertoires of both subgroups (excluding TAL effectors), from bacterial growth in barley leaves inoculated via spray at six days post inoculation, and from chlorophyll concentration in wheat leaves inoculated via infiltration at three days post inoculation were analyzed by Kruskal-Wallis test followed by Wilcoxon rank-sum test in R v4.1.0. Data from bacterial growth curves in planta upon syringe infiltration were analyzed per time point by one way analysis of variance (ANOVA) followed by Tukey’s post-hoc test in the log-transformed data. Assumptions were tested on the residuals of the model, and the normality of the data was assessed by the Shapiro-Wilk test in R v4.1.0 under the package rstatix (https://rpkgs.datanovia.com/rstatix/). Data from bacterial growth in barley leaves inoculated via spray at four days post inoculation were analyzed via ANOVA followed by Tukey’s post-hoc test upon checking normality of the data. The original data (x) of the percentage of secreted proteins uniquely found in pathovars translucens or undulosa, and of the percentage of symptomatic leaf area in barley leaves inoculated via spray were submitted to a transformation that combines the arcsine and square root functions (y = arcsin(√x/100)). The normality of data was then assessed by the Shapiro-Wilk test, and treatments were analyzed using the two-tailed Student’s *t*-test. Phylogenetic trees were visualized, and mid-point rooted via FigTree v1.4.4. (http://tree.bio.ed.ac.uk/). Genomic neighborhoods were visualized using Clinker.

## Acknowledgments

We thank Carine Gris (LIPME, Université de Toulouse, INRAE, CNRS) for the help in providing the *X. campestris* pv. campestris strains. We acknowledge funding from the Agriculture and Food Research Initiative SCRI grant no. 2020-51181-32154, from the INRAE SPE project XANTHOHS to E.L., L.M. and L.D.N., and from the American Malting Barley Association. M.V.M. is a President Post Doctoral Fellow of the President’s Postdoctoral Scholars Program of The Ohio State University. The authors declare that they have no competing interest.

