## Supplementary Tables and Figures for "Effector loss and gain drives pathogen host range at a fitness cost"

Marcus V. Merfa *et al.*

**This pdf file includes:**

Figs. S1 to S17.
Tables S3 to S4, and S6 to S9 (tables S1, S2, and S5 are provided as Microsoft Excel Files).
Supplementary Materials References.

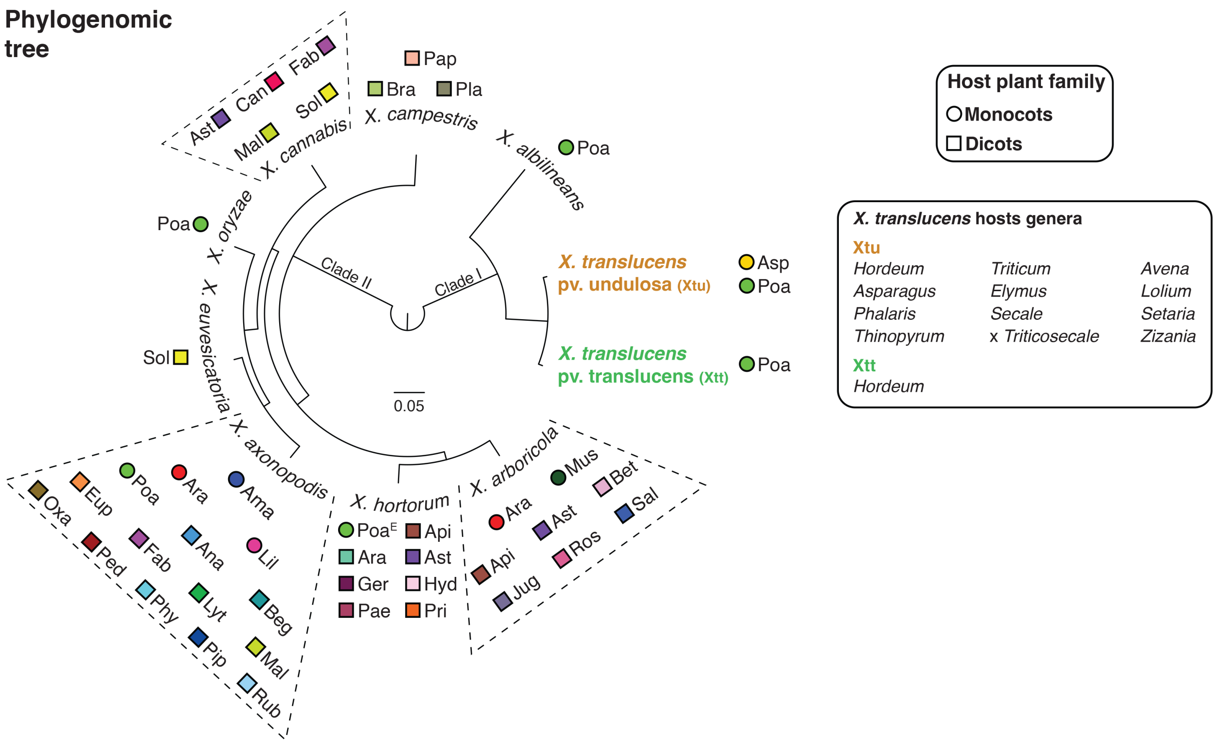

**Fig. S1.** Phylogenomics and host range of representative *Xanthomonas* spp. with differential host range breadth. The phylogenomic tree of representative *Xanthomonas* species with whole genome sequences was made from concatenated single-copy orthologous genes (SCOs). The tree was visualized via FigTree, and mid-point rooted. Plant families of natural hosts of each *Xanthomonas* spp. are color-coded and shown next to respective species in the tree (circle for monocots and square for dicots). Genera of host plants for *X. translucens* pv. translucens (*Hordeum*-only) and pv. undulosa are indicated in the figure. Monocots: Ama – *Amatyllidaceae*; Ara – *Araceae*; Asp – *Asparagaceae*; Lil – *Liliaceae*; Mus – *Musaceae*; Poa – *Poaceae*. Dicots: Ana – *Anacardiaceae*; Api – *Apiaceae*; Ara – *Araliceae*; Ast – *Asteraceae*; Beg – *Begoniaceae*; Bet – *Betulaceae*; Bra – *Brassicaceae*; Can – *Cannabaceae*; Eup – *Euphorbiaceae*; Fab – *Fabaceae*; Ger – *Geraniaceae*; Jug – *Juglandaceae*; Lyt – *Lythraceae*; Mal – *Malvaceae*; Oxa – *Oxalidaceae*; Pap – *Papaveraceae*; Ped – *Pedaliaceae*; Phy – *Phyllanthaceae*; Pip – *Piperaceae*; Pla – *Plantaginaceae*; Ros – *Rosaceae*; Rub – *Rubiaceae*; Sal – *Salicaceae*; Sol – *Solanaceae*. [E]Indicates experimentally determined host plant family.

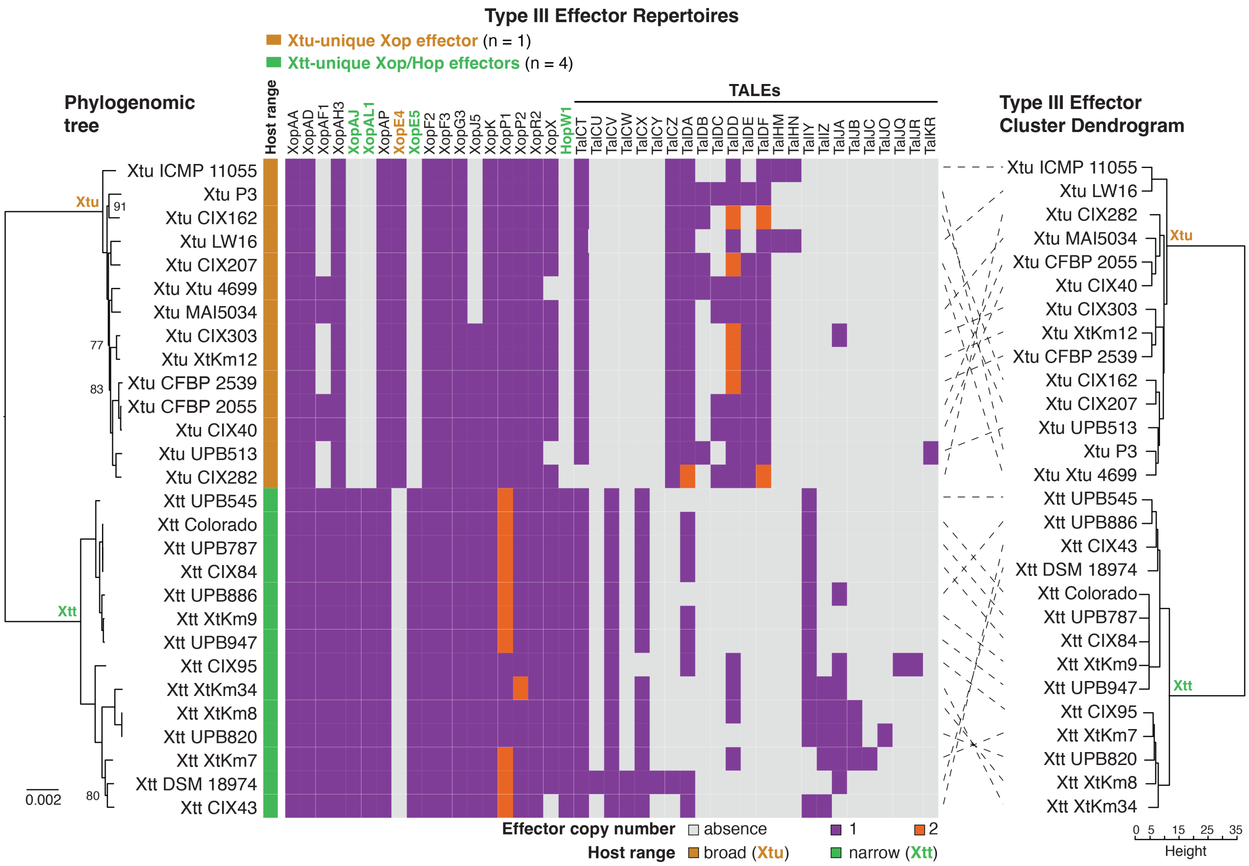

**Fig. S2.** Effector repertoires, including TALEs, of *Xanthomonas translucens* pathovars translucens, and undulosa. A core genome-based tree of 28 *X. translucens* strains is shown to the left side of the figure, demonstrating specific clustering of each pathovar. The tree was visualized via FigTree, and mid-point rooted. Branches with bootstrap values below 98% are indicated in the tree. Putative Type III secretion system effectors were identified via local Blastx against a database of known *Xanthomonas* effectors. Transcription activator-like effectors (TALEs) were identified and classified using AnnoTALE v.1.5. Inclusion of TALEs in the effector repertoire analysis disrupted the hierarchical clustering of strains within each pathovar according to their encoded repertoires (Type III effector cluster dendrogram; right side of the figure), making them not follow the phylogeny of the species (as shown in the phylogenomic tree; left side of the figure), contrary to what is obtained when TALEs are not included (see figure 1C). Xtt – *X. translucens* pv. translucens; Xtu – *X. translucens* pv. undulosa.

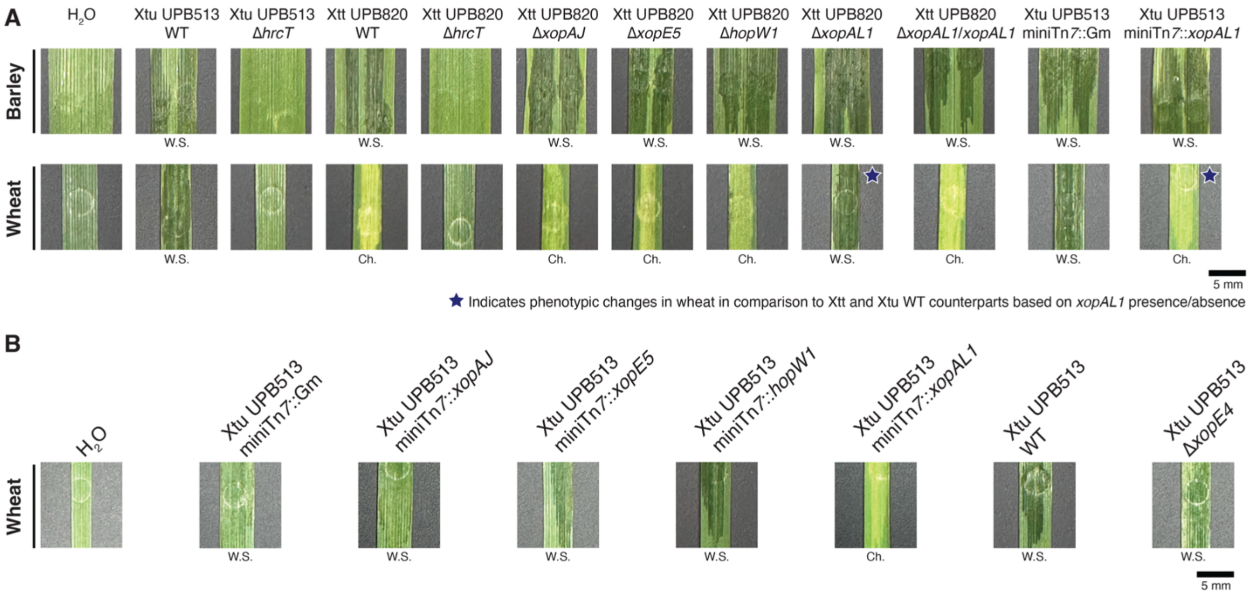

**Fig. S3.** XopAL1 is the only Xtt-unique T3E that allows for host jump from barley to wheat upon its deletion. **A.** Symptom development in barley and wheat leaves inoculated (syringe infiltration) with Xtu and Xtt strains (wild-type, mutant, and insertion strains; $OD_{600}$ 0.1) at three days post inoculation. Scale bar is shown in figure. **B.** XopAL1 is the only Xtt-unique T3E that triggers the immune response of wheat when introduced to Xtu UPB513 via mini-Tn*7* insertion; and deletion of the Xtu-unique effector *xopE4* does not affect its pathogenicity in wheat. Symptom development in wheat leaves inoculated (syringe infiltration) with Xtu strains ($OD_{600}$ 0.1) at three days post inoculation. Scale bar is shown in figure. Barley – *Hordeum vulgare* L. cv. Morex; Wheat – *Triticum aestivum* L. genotype 'Chinese Spring'; Xtt - *X. translucens* pv. translucens; Xtu - *X. translucens* pv. undulosa; Ch. – chlorosis; W.S. – water soaking.

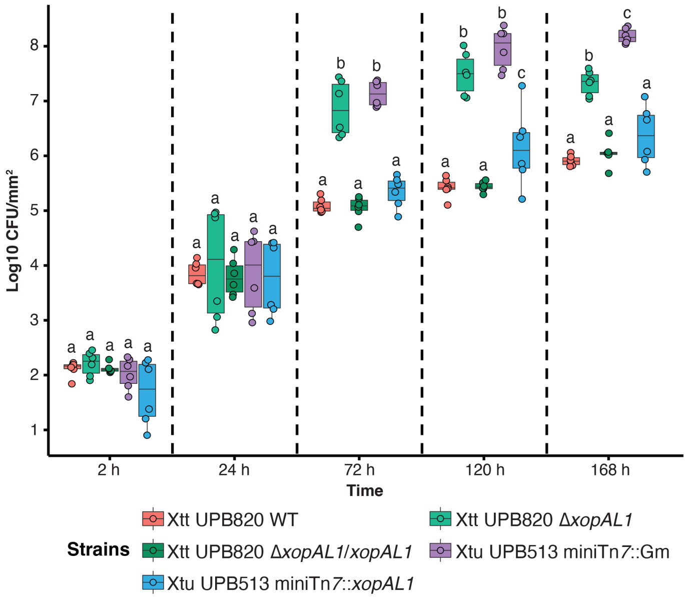

**Fig. S4.** Presence of *xopAL1* significantly reduces the population of *X. translucens* in wheat. Xtt and Xtu (wild-type, mutant, complemented and insertion strains for *xopAL1*) populations in planta determined at 24- to 48-hour intervals in wheat leaves. Xtt strains were previously transformed with pPneo-GFP, and Xtu strains were previously transformed via miniTn*7*-Gm (gentamicin) insertion, to allow for growth in selective medium. Leaves were inoculated with Xtt and Xtu strains via syringe infiltration ($OD_{600}$ 0.001). Different letters indicate significant difference per time point (separated by vertical dashed lines) as determined by one way ANOVA and Tukey's post-hoc test in the log-transformed data ($P<0.05$; n = two independent replicates with three biological replicates within each time point and inoculated strain, with three internal replicates each). Deletion of *xopAL1* in Xtt significantly increased the bacterial population in the non-host wheat, while expression of this effector gene in Xtu significantly reduced its population in wheat. Wheat – *Triticum aestivum* L. genotype 'Chinese Spring'; Xtt – *X. translucens* pv. translucens; Xtu – *X. translucens* pv. undulosa.

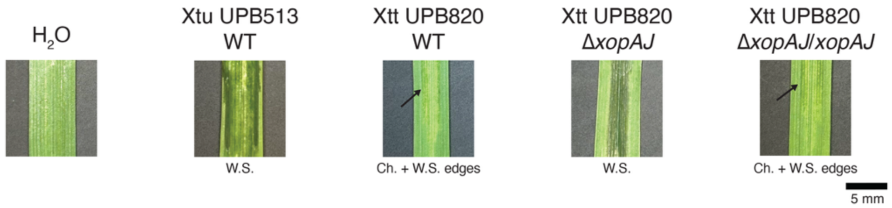

**Fig. S5.** Deletion of the T3E gene *xopAJ* allows for disease development caused by Xtt in oat. Symptom development in oat (*Avena sativa* L. variety Antigo) leaves inoculated (syringe infiltration) with Xtu and Xtt strains (wild-type, mutant, and complemented strains; $OD_{600}$ 0.1) at three days post inoculation. Arrows are pointing to water-soaked edges of chlorotic lesions. Scale bar is shown in figure. Xtt - *X. translucens* pv. translucens; Xtu - *X. translucens* pv. undulosa; Ch. – chlorosis; W.S. – water soaking.

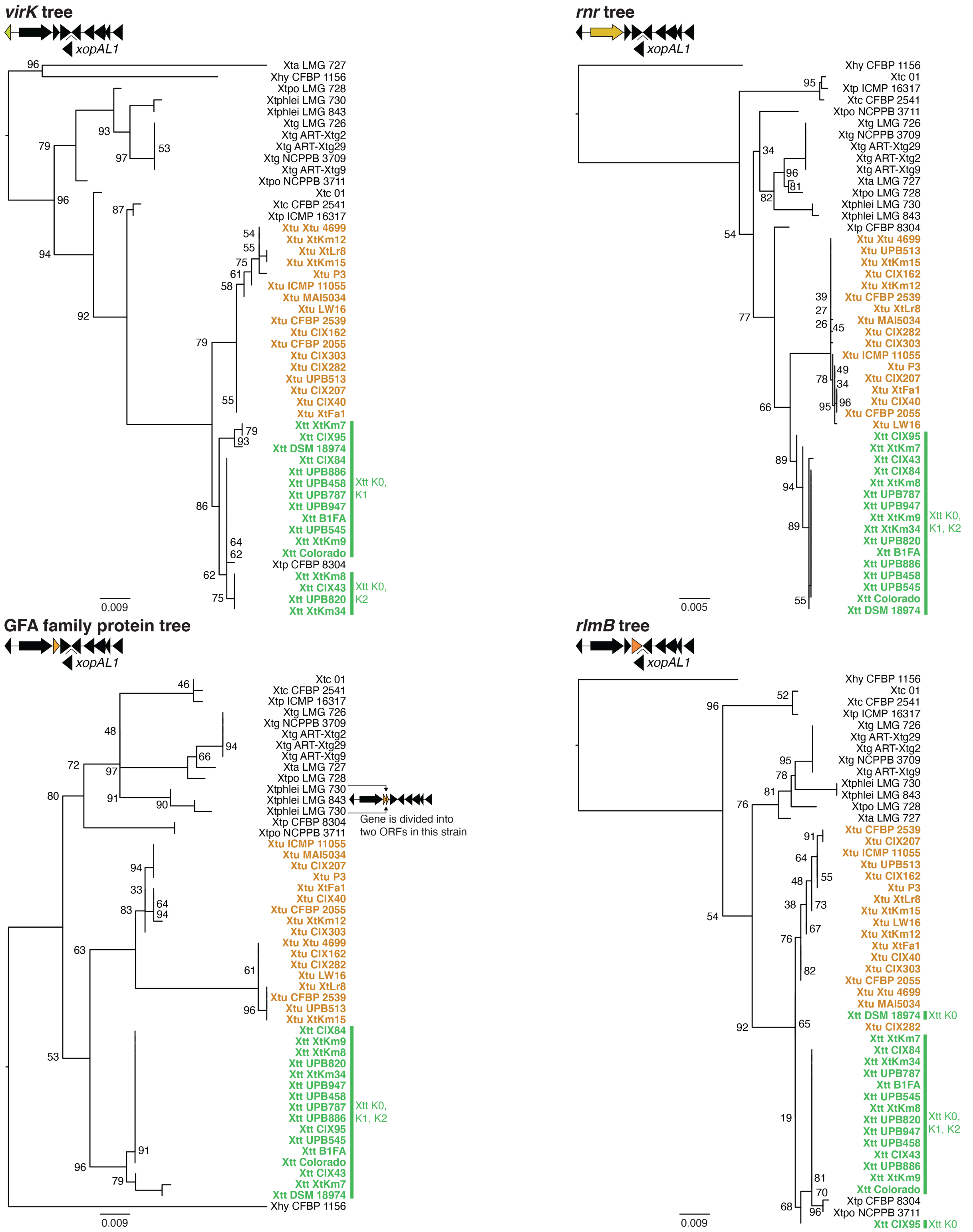

**Fig. S6.** Individual nucleotide-sequence-based phylogenetic trees of the four upstream genes within the *xopAL1* locus neighborhood of representative strains of *Xanthomonas translucens* pathovars with whole-genome sequences. *X. hyacinthi* strain CFBP 1156, a closely-related clade I *Xanthomonas* spp. to *X. translucens*, was included as an outgroup to root the trees. Orthologs of each gene were identified using OrthoFinder and trees were built using the Maximum-likelihood method via IQ-TREE. Trees were visualized via FigTree. Branches with bootstrap values below 98% are shown in each tree. The K0, K1 and K2 sub-genomic groups of *X. translucens* pv. translucens, as determined by life identification numbers (LINs) calculated with LINbase, are shown in the figure. Xhy – *X. hyacinthi*; Xta – *X. translucens* pv. arrhenateri; Xtpo – *X. translucens* pv. poae; Xtphlei – *X. translucens* pv. phleipratensis; Xtg – *X. translucens* pv. graminis; Xtc – *X. translucens* pv. cerealis; Xtp – *X. translucens* pv. pistaciae; Xtu – *X. translucens* pv. undulosa; Xtt – *X. translucens* pv. translucens.

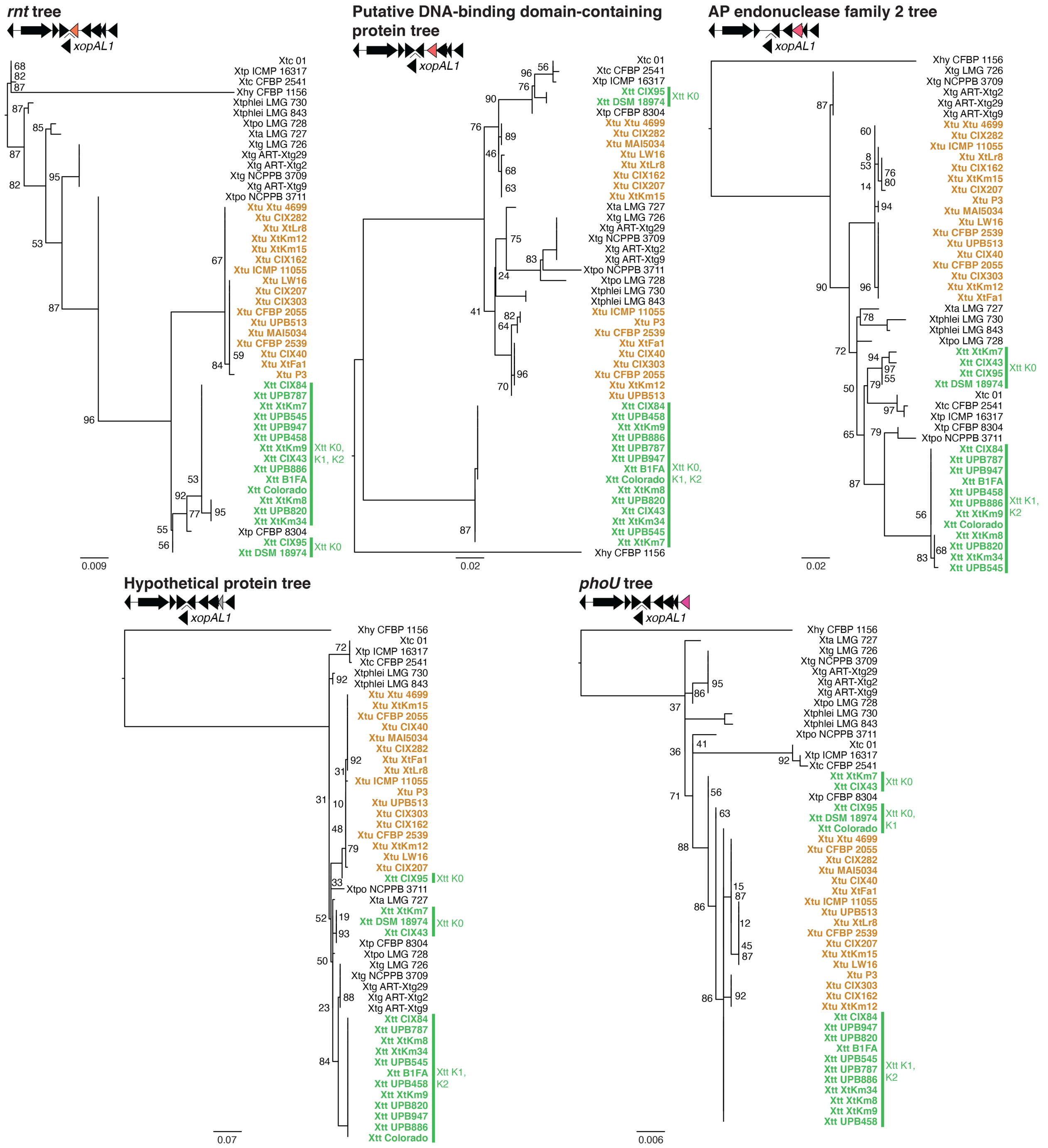

**Fig. S7.** Individual nucleotide-sequence-based phylogenetic trees of the five downstream genes within the *xopAL1* locus neighborhood of representative strains of *Xanthomonas translucens* pathovars with whole-genome sequences. *X. hyacinthi* strain CFBP 1156, a closely-related clade I *Xanthomonas* spp. to *X. translucens*, was included as an outgroup to root the trees. Orthologs of each gene were identified using OrthoFinder and trees were built using the Maximum-likelihood method via IQ-TREE. Trees were visualized via FigTree. Branches with bootstrap values below 98% are shown in each tree. The K0, K1 and K2 sub-genomic groups of *X. translucens* pv. translucens, as determined by life identification numbers (LINs) calculated with LINbase, are shown in the figure. Xhy – *X. hyacinthi*; Xtc – *X. translucens* pv. cerealis; Xtp – *X. translucens*

pv. pistaciae; Xtphlei – *X. translucens* pv. phleipratensis; Xtpo – *X. translucens* pv. poae; Xta – *X. translucens* pv. arrhenateri; Xtg – *X. translucens* pv. graminis; Xtu – *X. translucens* pv. undulosa; Xtt – *X. translucens* pv. translucens.

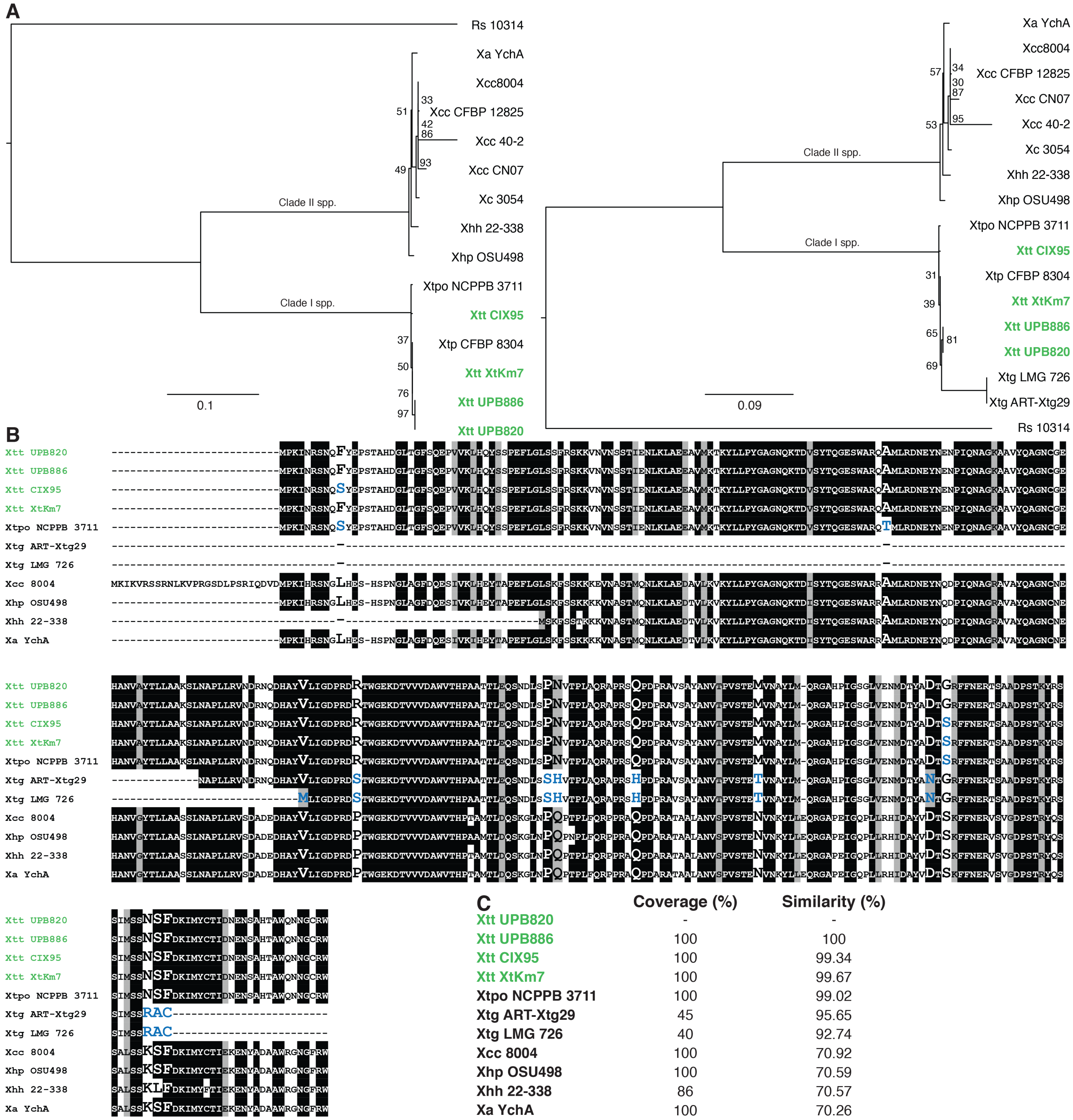

C

| | Coverage (%) | Similarity (%) |
|---|---|---|
| Xtt UPB820 | - | - |
| Xtt UPB886 | 100 | 100 |
| Xtt CIX95 | 100 | 99.34 |
| Xtt XtKm7 | 100 | 99.67 |
| Xtpo NCPPB 3711 | 100 | 99.02 |
| Xtg ART-Xtg29 | 45 | 95.65 |
| Xtg LMG 726 | 40 | 92.74 |
| Xcc 8004 | 100 | 70.92 |
| Xhp OSU498 | 100 | 70.59 |
| Xhh 22-338 | 86 | 70.57 |
| Xa YchA | 100 | 70.26 |

**Fig. S8.** XopAL1 diversity in *Xanthomonas* spp. **A.** Nucleotide-sequence-based phylogenetic trees of *xopAL1* sequences from representative clade I and clade II *Xanthomonas* spp. The tree to the left side of the panel does not contain the truncated *xopAL1* sequences that are encoded by strains of *X. translucens* pv. graminis, which are included in the tree to the right side of the panel. XopAL1 from *Ralstonia solanacearum* strain 10314 was included as an outgroup to root the tree. Trees were visualized via FigTree. Branches with bootstrap values below 98% are shown in each tree. **B.** Alignment of representative XopAL1 ortholog sequences within *Xanthomonas* spp. Sequences were aligned via T-Coffee and visualized using pyBoxShade. Different sequences of XopAL1 are denoted by representative strains within each species. Black shading indicates conserved residues; grey shading indicates conservative mutations; and white color indicates divergence among sequences. Amino acid changes within *X. translucens* is indicated by using a bigger font size and the color blue (in comparison to the reference XopAL1 encoded by Xtt UPB820). Sequences represent the entire diversity of XopAL1 within *X. translucens.* **C.** Percent coverage and similarity of XopAL1 sequences encoded among representative *Xanthomonas* spp. in comparison to the reference Xtt strain UPB820. Xa - *X. arboricola*; Xhh - *X. hortorum* pv. hederae; Xhp - *X. hortorum* pv. pelargonii; Xcc - *X. campestris* pv. campestris; Xc - *X. campestris*; Xtt - *X. translucens* pv. translucens; Xtp - *X. translucens* pv. pistaciae; Xtpo - *X. translucens* pv. poae; Xtg - *X. translucens* pv. graminis; Rs - *Ralstonia solanacearum*.

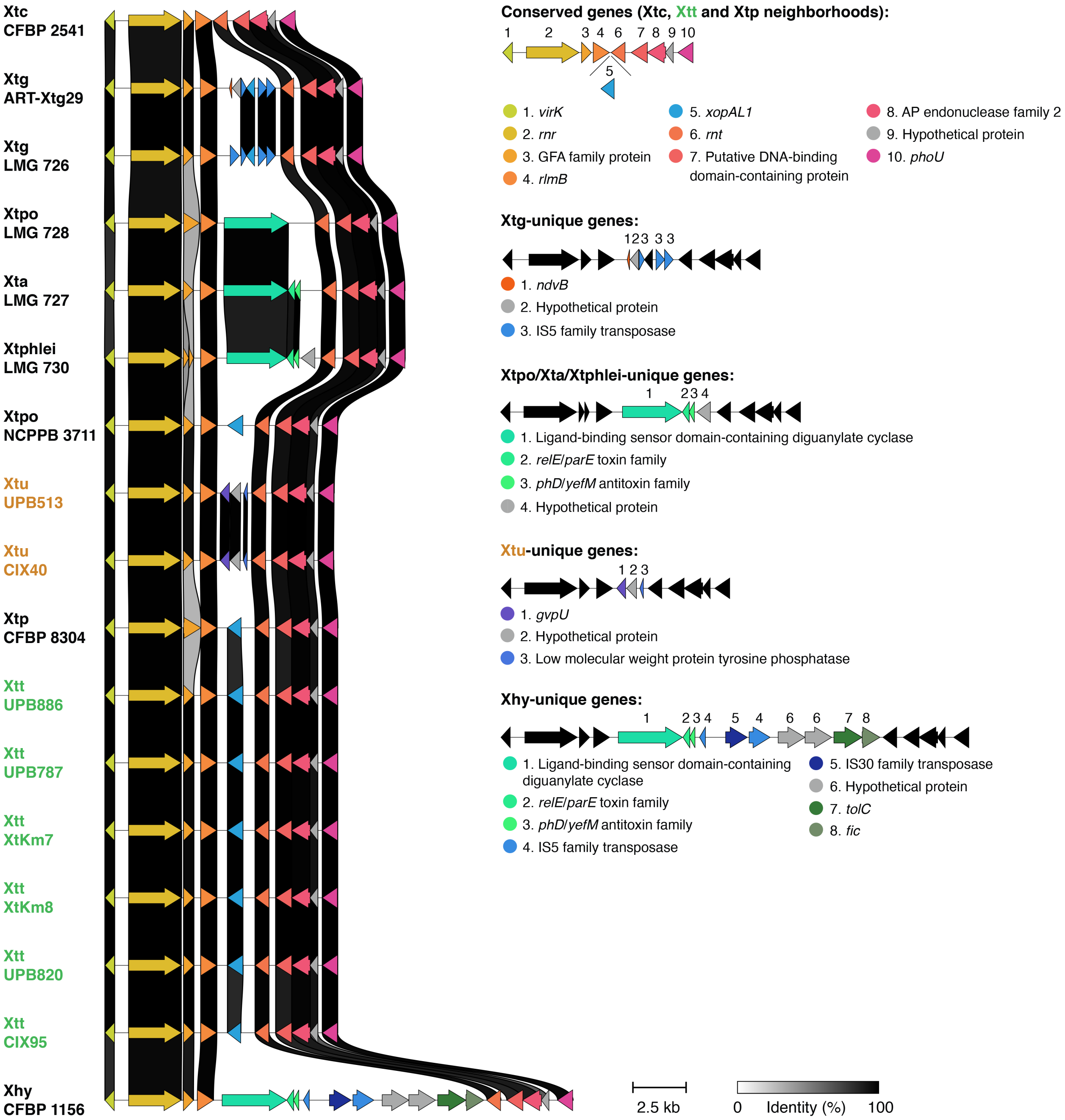

**Fig. S9.** Schematic overview of the *xopAL1* locus neighborhood types in representative pathovars of *Xanthomonas translucens* and in the closely related clade I species *Xanthomonas hyacinthi* using Clinker. Among analyzed strains, only Xtt, Xtp CFBP 8304 and Xtpo NCPPB 3711 encode *xopAL1*. Strains of Xtc, Xtt, Xtp CFBP8304 and Xtpo NCPPB 3711 have the same neighborhood, with Xtc sharing all genes except *xopAL1*. Xtg encodes truncated versions of *xopAL1*, which is likely due to the presence of IS5 family transposases. Within the other non-*xopAL1*-encoding strains, there is presence of other genes in place of *xopAL1*. Xtpo, Xta and Xtphlei strains have a similar neighborhood that shares similarities with the neighborhood of Xhy, which encodes additional genes in comparison to all pathovars of *X. translucens* (including transposases). At last, Xtu encodes a neighborhood that is specific to this pathovar. Xtc – *X. translucens* pv. cerealis; Xtg – *X. translucens* pv. graminis; Xtpo – *X. translucens* pv. poae; Xta

– *X. translucens* pv. arrhenateri; Xtphlei – *X. translucens* pv. phleipratensis; Xtu – *X. translucens* pv. undulosa; Xtp – *X. translucens* pv. pistaciae; Xtt – *X. translucens* pv. translucens; Xhy – *X. hyacinthi*.

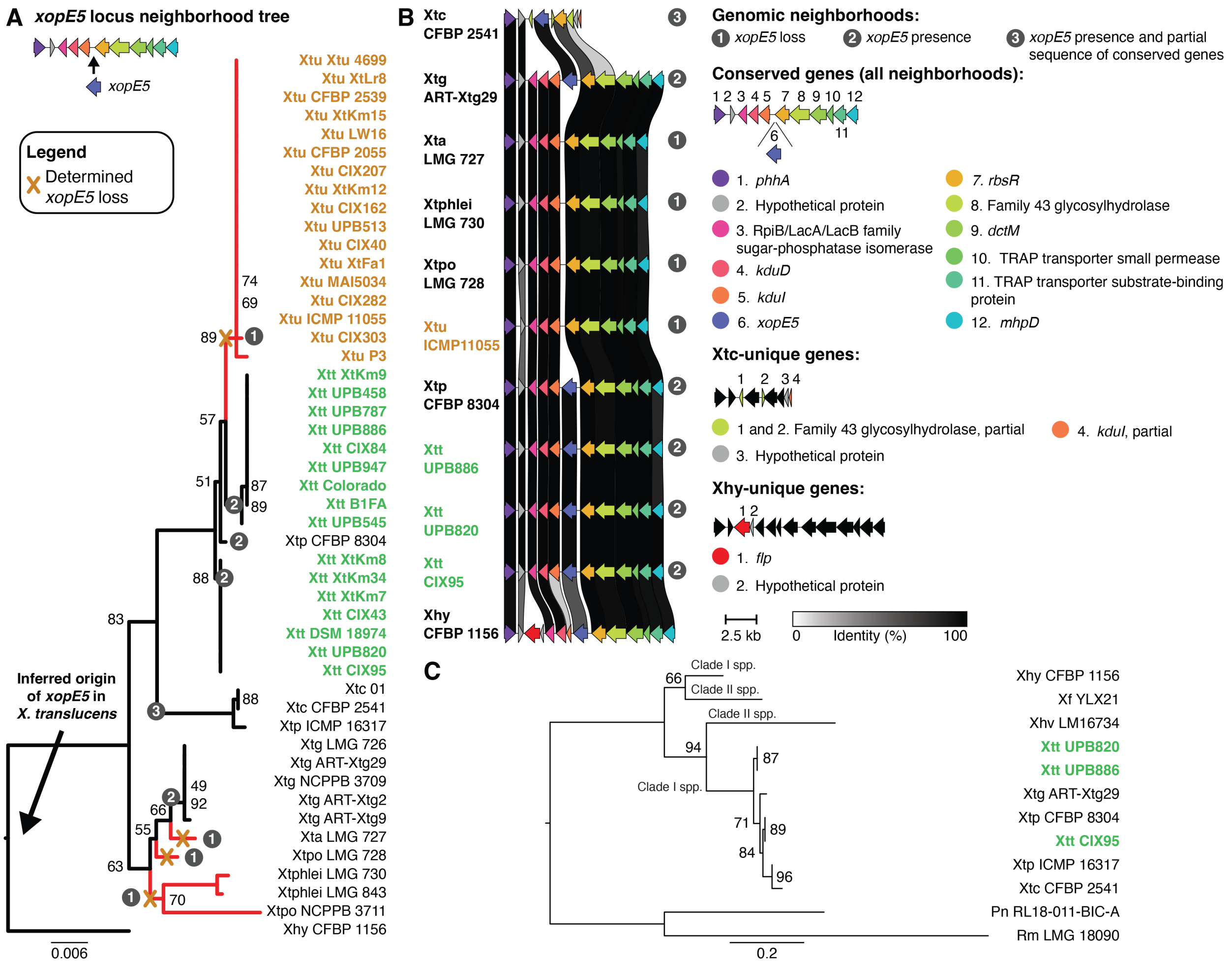

**Fig. S10.** *xopE5* has an ancestral origin in *Xanthomonas translucens* and was lost in Xtu. **A.** Tree of the concatenated sequences of the genes flanking *xopE5* in *X. translucens*. The *xopE5* locus neighborhood types are indicated in each branch and correspond to neighborhood types shown in panel B. Red-colored branches indicate subgroups that do not encode this effector gene, with determined loss events being shown (see tree legend). *X. hyacinthi*, a closely related clade I *Xanthomonas* spp., was included as an outgroup to root the tree. **B.** Schematic overview of the *xopE5* locus neighborhood types in representative pathovars of *Xanthomonas translucens* and in the closely related species *X. hyacinthi* using Clinker. Among analyzed strains, all Xtt, Xtp, Xtc, and *X. hyacinthi* encode *xopE5*. All these have similar neighborhoods. However, Xtc strains have a smaller neighborhood with partial sequence of conserved genes found in all strains. Within non-*xopE5*-encoding strains, there are no other genes replacing *xopE5*. *X. hyacinthi* encodes two additional genes in comparison to all pathovars of *X. translucens*. **C.** Nucleotide-sequence-based phylogenetic tree of *xopE5* sequences from representative clade I and clade II *Xanthomonas* spp. *xopE5* sequences from *Paraburkholderia nemoris* strain RL18-011-BIC-A and *Ralstonia mannitolilytica* strain LMG 18090 were used as outgroups to root the tree. *xopE5* is encoded in a similar neighborhood by the closely related species *X. hyacinthi* (panel B), and this ortholog is closely related to *xopE5* encoded by *X. translucens* pathovars (panel C). Thus, we inferred an ancestral origin of this effector gene in *X. translucens* (i.e. common origin with a shared ancestor with *X. hyacinthi*), and determined subsequent evolutionary losses in *X. translucens* subgroups that do not encode this gene. This includes loss of *xopE5* in Xtu during its divergence from the common ancestor with Xtt. Trees were visualized via Figtree. Branches with bootstrap values below 98% are indicated in each tree. Xtu – *X. translucens* pv. undulosa; Xtt – *X. translucens* pv. translucens; Xtp – *X. translucens* pv. pistaciae; Xtc – *X. translucens* pv. cerealis; Xtg – *X. translucens* pv. graminis; Xta – *X. translucens* pv. arrhenateri; Xtpo – *X. translucens* pv. poae; Xtphlei – *X. translucens* pv. phleipratensis; Xhy – *X. hyacinthi*; Xf – *X. fragariae*; Xhv – *X. hortorum* pv. vitians; Pn – *Paraburkholderia nemoris*; Rm – *Ralstonia mannitolilytica*.

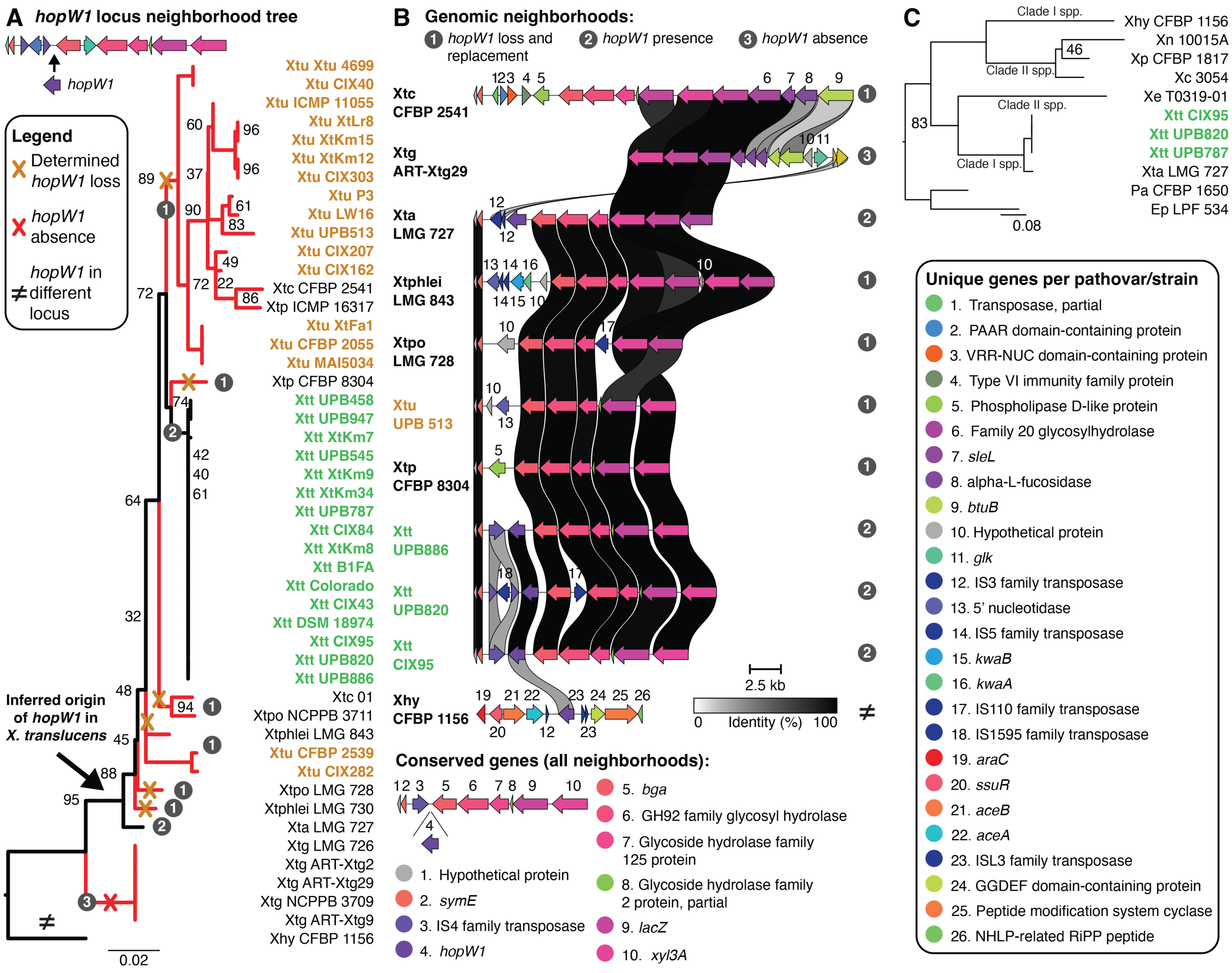

**Fig. S11.** *hopW1* was lost in Xtu and is encoded in a highly rearranged genomic neighborhood. **A.** Tree of the concatenated sequences of the genes flanking *hopW1* in *X. translucens*. The *hopW1* locus neighborhood types are indicated in each branch and correspond to neighborhood types shown in panel B. Red-colored branches indicate subgroups that do not encode this effector gene, with determined absence and loss events being shown (see tree legend). *X. hyacinthi*, a closely related clade I *Xanthomonas* spp., was included as an outgroup to root the tree. **B.** Schematic overview of the *hopW1* locus neighborhood types in representative pathovars of *Xanthomonas translucens* and in the closely related species *X. hyacinthi* using Clinker. Among analyzed strains, all Xtt, Xta LMG 727, and *X. hyacinthi* encode *hopW1*. Xtt strains encode *hopW1* in similar genomic neighborhoods, but which have few differences in comparison to the neighborhood of Xta LMG 727. The latter encodes two additional genes (IS3 family transposases) in comparison to Xtt strains. *X. hyacinthi* encodes *hopW1* in a different genomic neighborhood in comparison to *X. translucens*. Within non-*hopW1*-encoding strains, there is presence of other genes in place of *hopW1*. **C.** Nucleotide-sequence-based phylogenetic tree of *hopW1* sequences from representative clade I and clade II *Xanthomonas* spp. *hopW1* sequences from *Pseudomonas amygdali* strain CFBP 1650 and *Erwinia psidii* strain LPF 534 were used as outgroups to root the tree. *hopW1* is encoded in a different neighborhood by the closely related species *X. hyacinthi* (panel B), and this ortholog is distantly related to *hopW1* encoded by *X. translucens* pathovars (panel C). Since the Xta and Xtt orthologs are closely related (panel C) and they share a similar genomic neighborhood (panel B), we inferred an origin in Xta upon divergence from Xtg, and subsequent losses in pathovars that do not encode this effector gene. This includes loss of *hopW1* in Xtu during its divergence from the common ancestor with Xtt. In addition, we observed rearrangements and transposases within this genomic neighborhood in *X. translucens* (see panel B), suggesting that this locus is subject to dynamic evolutionary events. Trees were visualized via Figtree. Branches with bootstrap values below 98% are indicated in each tree. Xtu – *X. translucens* pv. undulosa; Xtc – *X. translucens* pv. cerealis; Xtp – *X. translucens* pv. pistaciae; Xtt – *X. translucens* pv. translucens; Xtpo – *X. translucens* pv. poae; Xtphlei – *X. translucens* pv. phleipratensis; Xta – *X. translucens* pv. arrhenateri; Xtg – *X. translucens* pv. graminis; Xhy – *X. hyacinthi*; Xn – *X. nasturtii*; Xp – *X. populi*; Xc – *X. campestris*; Xe – *X. euvesicatoria*; Pa – *Pseudomonas amygdali*; Ep – *Erwinia psidii*.

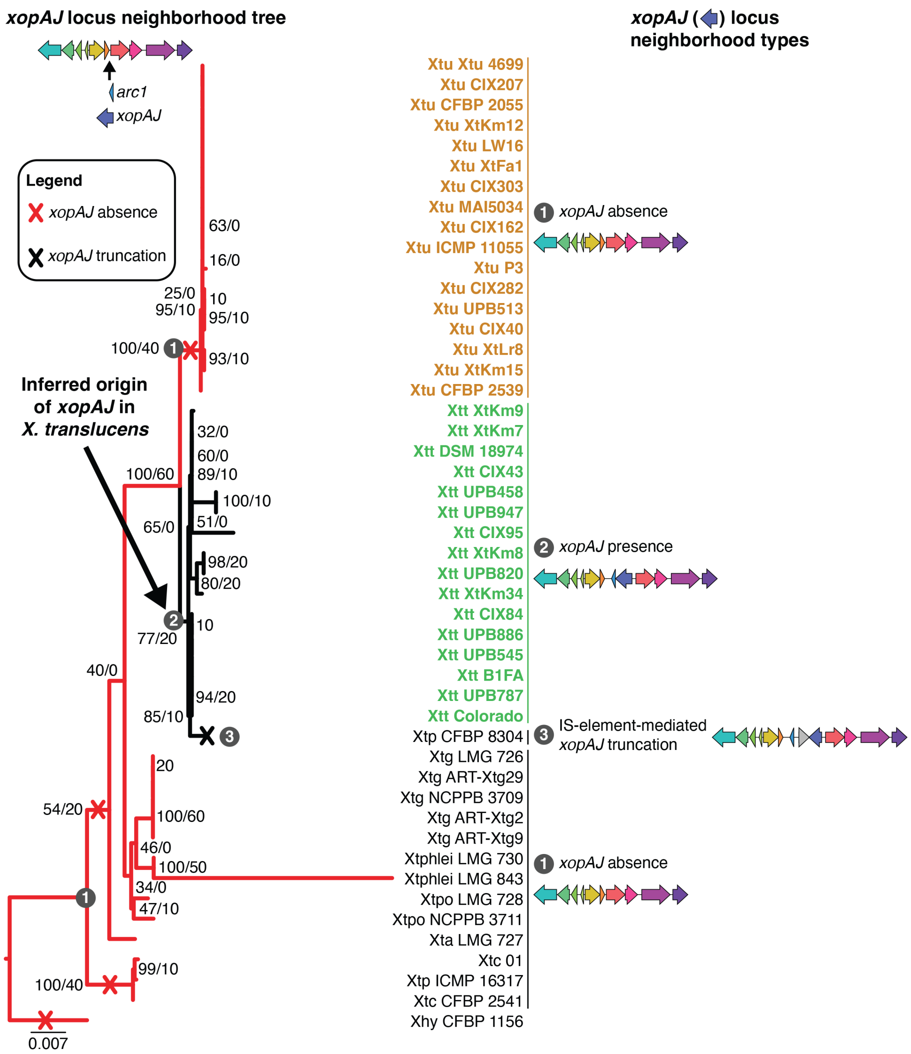

**Fig. S12.** *xopAJ* originated in *X. translucens* pv. translucens after divergence with *X. translucens* pv. undulosa and was truncated in pathovar pistaciae strain CFBP 8304. Tree of the concatenated amino acid sequences of the genes flanking *xopAJ* (6 upstream and 4 downstream genes) in *X. translucens*. The *xopAJ* locus neighborhood types are highlighted within each branch and shown to the right side of the figure (see Fig. S13 for more details). Red-colored branches indicate subgroups that do not encode this effector, with absence being shown (see tree legend). *X. hyacinthi* strain CFBP 1156, a closely-related clade I *Xanthomonas* spp., was included as an outgroup to root the tree. The tree was visualized via Figtree. Branches are labeled with bootstrap values followed by the gene concordance factor (gCF). Xtu – *X. translucens* pv. undulosa; Xtt – *X. translucens* pv. translucens; Xtp – *X. translucens* pv. pistaciae; Xtc – *X. translucens* pv. cerealis; Xtg – *X. translucens* pv. graminis; Xtphlei – *X. translucens* pv. phleipratensis; Xtpo – *X. translucens* pv. poae; Xta – *X. translucens* pv. arrhenateri; Xhy – *X. hyacinthi*.

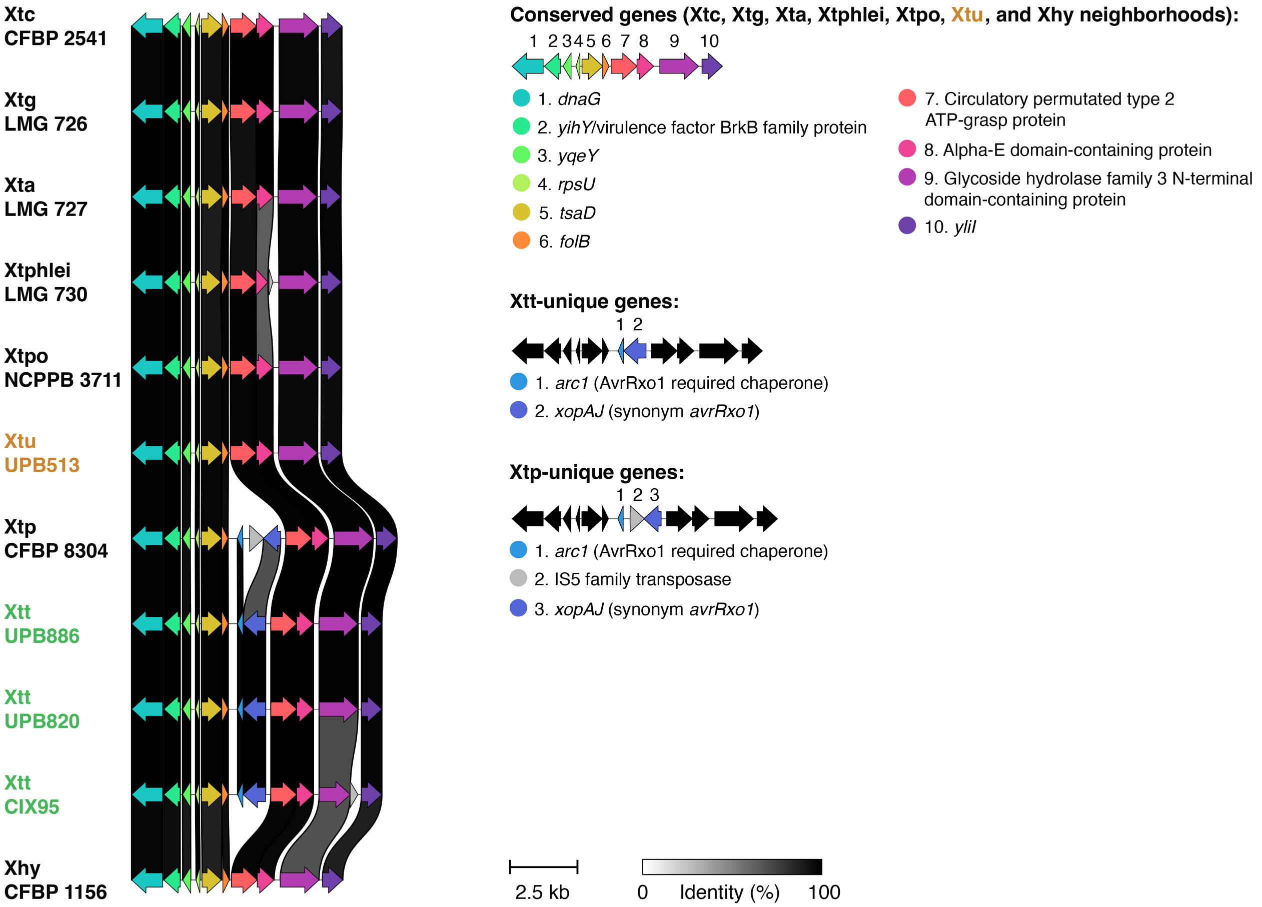

**Fig. S13.** Schematic overview of the *xopAJ* locus neighborhood types in representative pathovars of *Xanthomonas translucens* and in the closely related clade I species *Xanthomonas hyacinthi* using Clinker. Among analyzed strains, only Xtt encodes *xopAJ*. Xtp CFBP8304 encodes a truncated version of *xopAJ*, which is likely due to the presence of an IS5 family transposase. Within non-*xopAJ*-encoding strains, there are no other genes replacing *xopAJ*. Xtc – *X. translucens* pv. cerealis; Xtg – *X. translucens* pv. graminis; Xta – *X. translucens* pv. arrhenateri; Xtphlei – *X. translucens* pv. phleipratensis; Xtpo – *X. translucens* pv. poae; Xtu – *X. translucens* pv. undulosa; Xtp – *X. translucens* pv. pistaciae; Xtt – *X. translucens* pv. translucens; Xhy – *X. hyacinthi*.

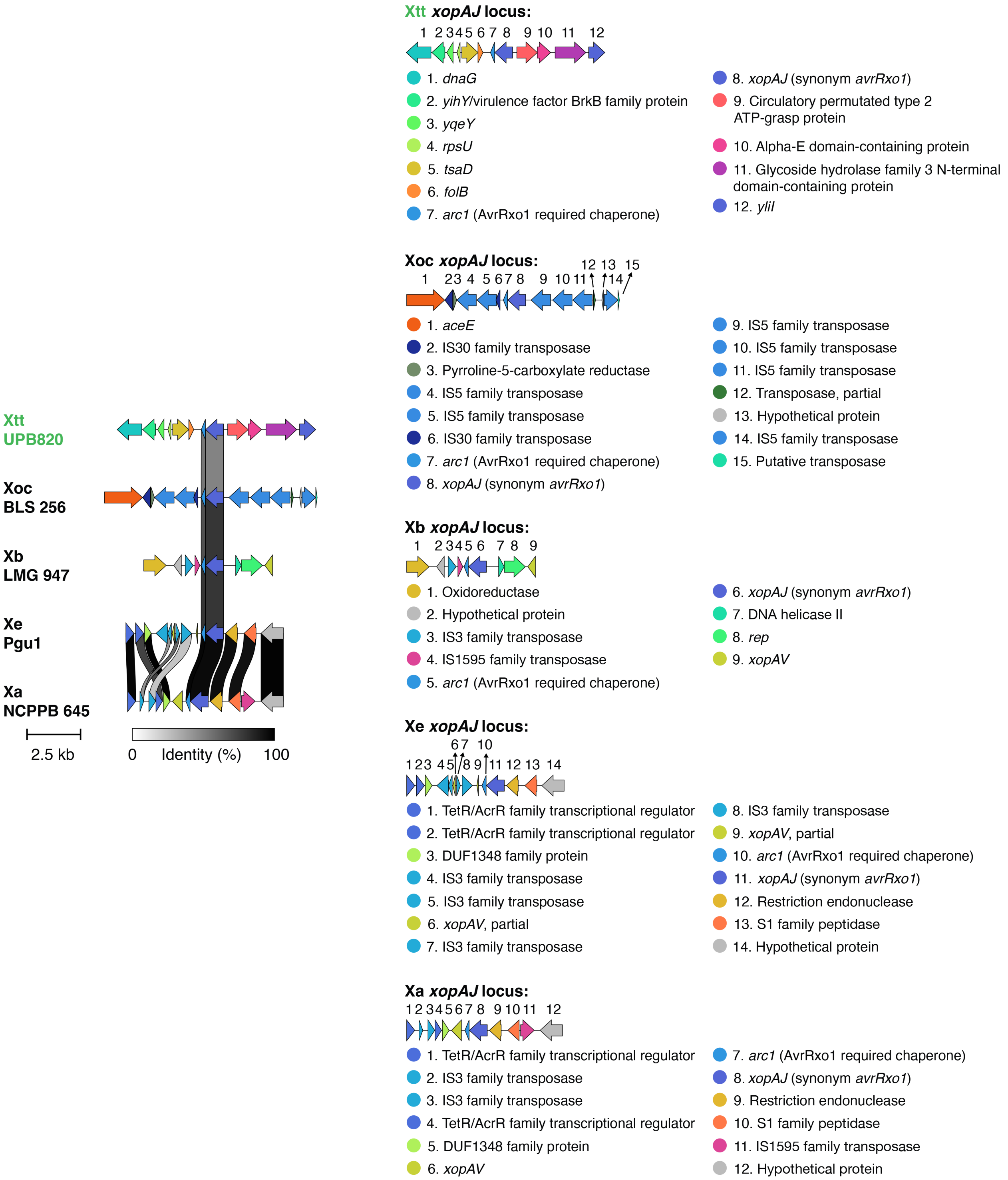

**Fig. S14.** Schematic overview of the *xopAJ* locus neighborhood types in Xtt UPB820 and representative clade II *Xanthomonas* species using Clinker. The *xopAJ* locus neighborhood is different between Xtt (clade I *Xanthomonas* spp.) and clade II *Xanthomonas* spp. Xoc and Xb have distinct *xopAJ* locus neighborhood types between themselves and in comparison to other clade II *Xanthomonas* spp. Xe and Xa have similar neighborhood types. All clade II *Xanthomonas* spp. have multiple insertion sequence (IS) elements in their *xopAJ* locus neighborhood types. Xtt – *X. translucens* pv. translucens; Xoc – *X. oryzae* pv. oryzicola; Xb – *X. bromi*; Xe – *X. euvesicatoria*; *Xa* – *X. axonopodis*.

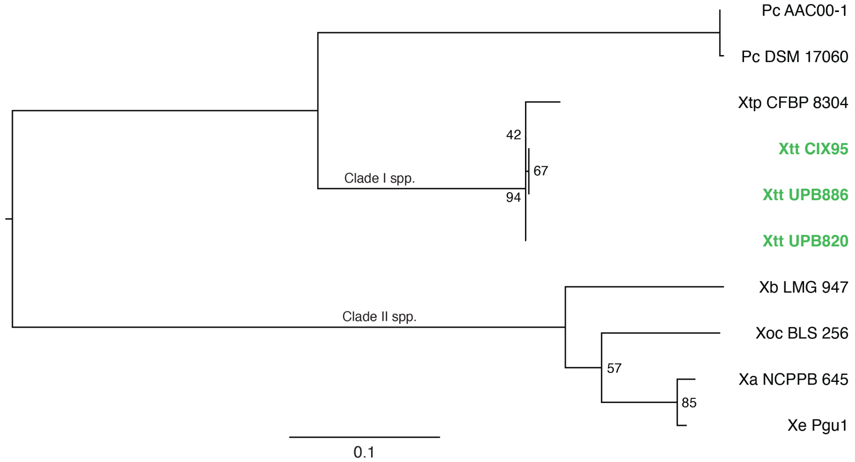

**Fig. S15.** Nucleotide-sequence-based phylogenetic tree of *xopAJ* sequences from representative clade I and clade II *Xanthomonas* spp. The tree was visualized via FigTree, and mid-point rooted. Branches with bootstrap values below 98% are shown in the tree. *xopAJ* sequences from *Paracidovorax citrulli* strains AAC00-1 and DSM 17060 were included as outgroups but clustered with *xopAJ* encoded by clade I *Xanthomonas* spp. Pc – *Paracidovorax citrulli*; Xtp – *X. translucens* pv. pistaciae; Xtt – *X. translucens* pv. translucens; Xb – *X. bromi*; Xoc – *X. oryzae* pv. oryzicola; *Xa* – *X. axonopodis*; Xe – *X. euvesicatoria*.

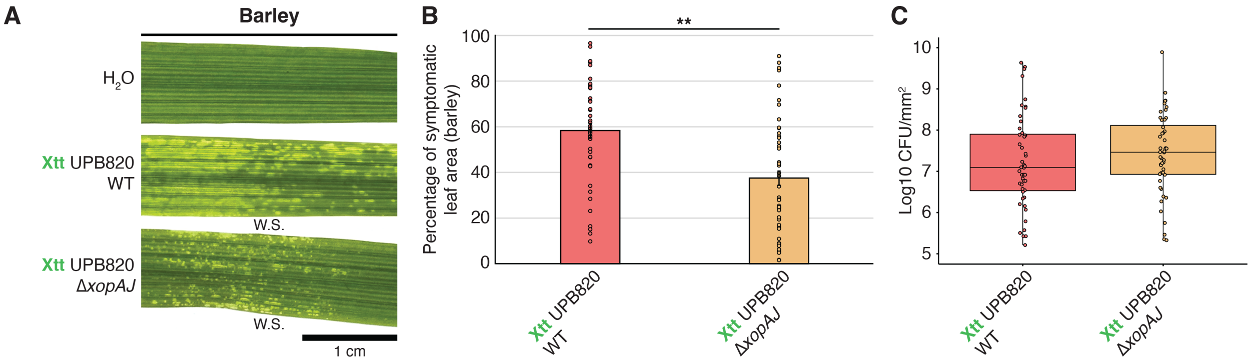

**Fig. S16.** Deletion of the T3E gene *xopAJ* decreases virulence of Xtt in barley. **A.** Symptom development in barley leaves inoculated (spray) with Xtt WT and Xtt Δ*xopAJ* ($OD_{600}$ 0.5) at four days post inoculation. Water soaking symptoms were visualized using a light box and appear as lighter areas in the leaf. Scale bar is shown in figure. **B.** Percentage of symptomatic leaf area of inoculated barley leaves (spray) with Xtt WT and Xtt Δ*xopAJ* shown in panel A. Total leaf area and symptomatic leaf area were calculated using ImageJ. Original data of percentage of symptomatic leaf area (treatments; y-axis) was submitted to a transformation that combines the arcsine and square root functions. The Shapiro-Wilk test (W) was then applied to the transformed data, and it supported the assumption of normal distribution (W = 0.982; *P* value = 0.223). ** indicates significant difference as determined by Student's *t*-test ($P<0.001$; n = two independent replicates with 24 biological replicates each). **C.** Xtt WT and Xtt Δ*xopAJ* populations in barley leaves inoculated via spray ($OD_{600}$ 0.5) at four days post inoculation. No significant differences between endophytic populations of Xtt WT and Xtt Δ*xopAJ* were observed according to one-way ANOVA and Tukey's post-hoc test ($P>0.05$; n = two independent replicates with 24 biological replicates each). Barley – *Hordeum vulgare* L. cv. Morex; Xtt - *X. translucens* pv. translucens; W.S. – water soaking.

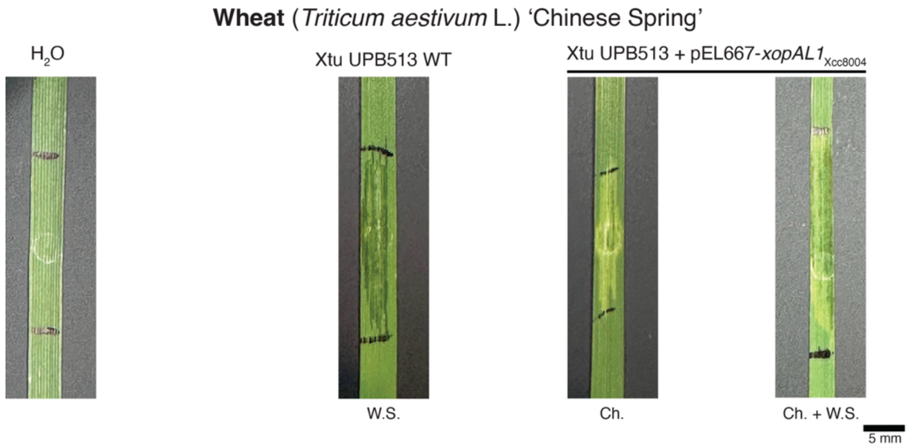

**Fig. S17.** The *X. campestris* pv. campestris (Xcc) strain 8004 XopAL1 ortholog also triggers wheat non-host response. Symptom development in wheat (*Triticum aestivum* L. genotype 'Chinese Spring') leaves inoculated (syringe infiltration) with Xtu strains ($OD_{600}$ 0.1) at three days post inoculation. The Xcc 8004 XopAL1 ortholog was expressed in Xtu via a plasmid construct, leading to development of chlorosis or a mix of chlorosis and water soaking when inoculated into wheat leaves. Scale bar is shown in figure. Xtu – *X. translucens* pv. undulosa; Ch. – chlorosis; W.S. – water soaking.

**Table S3.** Symptom development assessment in a cereal genotype diversity panel inoculated with Xtt and Xtu strains.

| | | | | Xanthomonas translucens strain | | | | |
|---|---|---|---|---|---|---|---|---|
| | | | | **Xtt UPB820 WT** | **Xtt UPB820 ∆*xopAL1*** | **Xtt UPB820 ∆*xopAL1*/∆*xopAL1*** | **Xtu UPB513 miniTn7::Gm** | **Xtu UPB513 miniTn7::*xopAL1*** |
| **Host** | **Scientific name** | **Genotype** | **Cultivar/variety/ genotype/line** | **Symptom** | | | | |
| Wheat | *Triticum aestivum* L. | BBAADD (2n=6x) | Chinese Spring[a] | Ch. | W.S. | Ch. | W.S. | Ch. |
| Wheat | *Triticum aestivum* L. | BBAADD (2n=6x) | RB07[a] | Ch. | W.S. | Ch. | W.S. | Ch. |
| Wheat | *Triticum aestivum* L. | BBAADD (2n=6x) | Green Spring[a] | Ch. | W.S. | Ch. | W.S. | Ch. |
| Wheat | *Triticum aestivum* L. | BBAADD (2n=6x) | AGS 2055[b] | Ch. | W.S. | Ch. | W.S. | Ch. |
| Wheat | *Triticum aestivum* L. *compactum* (Host) MacKey | BBAADD (2n=6x) | Cara[c] | W.S. + Ch. | W.S. | N.A. | W.S. | N.A. |
| Durum Wheat | *Triticum turgidum* L. ssp. *durum* (Desf.) Husnot | BBAA (2n=4x) | Lustre[d] | W.S. | W.S. | Ch. / W.S. | W.S. | W.S. |
| Durum Wheat | *Triticum turgidum* L. ssp. *durum* (Desf.) Husnot | BBAA (2n=4x) | ND Riveland[d] | Ch. / W.S. | W.S. | Ch. / W.S. | W.S. | Ch. / W.S. |
| Einkorn Wheat | *Triticum monococcum* ssp. *boeoticum* | AA (2n) | - | Ch. / W.S. | Ch. / W.S. | Ch. / W.S. | W.S. | W.S. |
| Emmer Wheat | *Triticum turgidum* ssp. *dicoccum* | BBAA (2n=4x) | - | W.S. | W.S. | W.S. | W.S. | W.S. |
| Barley | *Hordeum vulgare* L. | (2n) | Morex[e] | W.S. | W.S. | W.S. | W.S. | W.S. |
| Barley | *Hordeum vulgare* L. | (2n) | Kewaunee[e] | W.S. | W.S. | W.S. | W.S. | W.S. |
| Barley | *Hordeum vulgare* L. | (2n) | Conlon[f] | W.S. | W.S. | W.S. | W.S. | W.S. |
| Barley | *Hordeum vulgare* L. | (2n) | ND Pinnacle[f] | W.S. | W.S. | W.S. | W.S. | W.S. |
| Rye (Spring) | *Secale cereale* L. | (2n) | - | Ch. | Ch. | Ch. | W.S. | W.S. / W.S. + Ch. |
| Rye (Winter) | *Secale cereale* L. | (2n) | - | Ch. | Ch. | Ch. | W.S. | W.S. |
| Rye | *Secale cereale* L. | (2n) | AC Remington[g] | Ch. / W.S. + Ch. | Ch. | Ch. / W.S. + Ch. | W.S. | W.S. |
| Triticale | *x Triticosecale Wittmack* | BBAARR (2n=6x) | - | Ch. | W.S. | Ch. | W.S. | W.S. |
| Triticale | *x Triticosecale Wittmack* | BBAARR (2n=6x) | - | Ch. / W.S. + Ch. | W.S. | Ch. / W.S. + Ch. | W.S. | W.S. |
| Triticale | *x Triticosecale Wittmack* | BBAARR (2n=6x) | H | Ch. | W.S. | Ch. | Ch. / W.S. | W.S. |
| Oat | *Avena sativa L.* | AACCDD (2n=6x) | Antigo[h] | Ch. + W.S. edges | Ch. + W.S. edges | N.A. | W.S. | W.S. |
| Intermediate Wheatgrass | *Thinopyrum intermedium* | $E^eE^eE^bE^bStSt$ (2n=6x) | - | Ch. | Ch. | Ch. | W.S. | W.S. |

Leaf inoculations were performed by syringe infiltration ($OD_{600}$ of 0.1) using two-week-old plants, and symptoms were visualized at three days post inoculation.
[a]Hard Red Spring Wheat; [b]Soft Red Winter Wheat; [c]Soft White Winter Wheat; [d]Spring Durum Wheat; [e]Six-row Spring Barley; [f]Two-row Spring Barley; [g]Winter Rye; [h]Spring oat. Ch. – chlorosis (non-host response); W.S. – water soaking (symptom development – susceptibility); N.A. – not analyzed. + indicates both symptoms present in the same inoculated leaf area; / indicates symptoms identified in different inoculated leaf areas.

**Table S4.** Symptom development assessment in a subset of the cereal genotype diversity panel inoculated with Xtt and Xtu strains.

| | | | | Xanthomonas translucens strain | | | | | |
|---|---|---|---|---|---|---|---|---|---|
| | | | | **Xtt UPB820 WT** | **Xtt UPB820 Δ*xopAJ*** | **Xtt UPB820 Δ*xopAJ*/Δ*xopAJ*** | **Xtt UPB820 Δ*xopE5*** | **Xtt UPB820 ΔhopW1** | **Xtu UPB513 WT** |
| **Host** | **Scientific name** | **Genotype** | **Cultivar/variety/ genotype/line** | **Symptom** | | | | | |
| Wheat | *Triticum aestivum* L. | BBAADD (2n=6x) | AGS 2055[a] | Ch. | Ch. | N.A. | Ch. | Ch. | W.S. |
| Durum Wheat | *Triticum turgidum* L. ssp. *durum* (Desf.) Husnot | BBAA (2n=4x) | Lustre[b] | W.S. | W.S. | N.A. | W.S. | W.S. | W.S. |
| Einkorn Wheat | *Triticum monococcum* ssp. *boeoticum* | AA (2n) | - | Ch. + W.S. spots | Ch. | N.A. | Ch. + W.S. spots | Ch. + W.S. spots | W.S. |
| Emmer Wheat | *Triticum turgidum* ssp. *dicoccum* | BBAA (2n=4x) | - | W.S. | W.S. | N.A. | W.S. | W.S. | W.S. |
| Barley | *Hordeum vulgare* L. | (2n) | Morex[c] | W.S. | W.S. | N.A. | W.S. | W.S. | W.S. |
| Barley | *Hordeum vulgare* L. | (2n) | Conlon[d] | W.S. | W.S. | N.A. | W.S. | W.S. | W.S. |
| Rye (Spring) | *Secale cereale* L. | (2n) | - | Ch. | Ch. | N.A. | Ch. | Ch. | W.S. |
| Rye (Winter) | *Secale cereale* L. | (2n) | - | Ch. | Ch. | N.A. | Ch. | Ch. | W.S. |
| Triticale | *x Triticosecale Wittmack* | BBAARR (2n=6x) | - | Ch. | Ch. | N.A. | Ch. | Ch. | W.S. |
| Oat | *Avena sativa L.* | AACCDD (2n=6x) | Antigo[g] | Ch. + W.S. edges | W.S. | Ch. + W.S. edges | Ch. + W.S. edges | Ch. + W.S. edges | W.S. |
| Intermediate Wheatgrass | *Thinopyrum intermedium* | $E^eE^eE^bE^b$StSt (2n=6x) | - | Ch. | Ch. | N.A. | Ch. | Ch. | W.S. |

Leaf inoculations were performed by syringe infiltration ($OD_{600}$ of 0.1) using two-week-old plants, and symptoms were visualized at three days post inoculation.
[a]Soft Red Winter Wheat; [b]Spring Durum Wheat; [c]Six-row Spring Barley; [d]Two-row Spring Barley; [g]Spring oat.
Ch. – chlorosis (non-host response); W.S. – water soaking (symptom development – susceptibility); N.A. – not analyzed. + indicates both symptoms present in the same inoculated leaf area; / indicates symptoms identified in different inoculated leaf areas.

**Table S6.** Bacterial strains and plasmids used in this study.

| Bacterial strain | Genotype or description | Source |
|---|---|---|
| ***Xanthomonas translucens* pv. translucens (Xtt)** | | |
| UPB820 WT | Wild-type Xtt strain UPB820 | [1,2] |
| UPB820 Δ*hrcT* | UPB820 with chromosomal *hrcT* deletion | This study |
| UPB820 Δ*xopAJ* | UPB820 with chromosomal *xopAJ* deletion | This study |
| UPB820 Δ*xopAL1* | UPB820 with chromosomal *xopAL1* deletion | This study |
| UPB820 Δ*xopE5* | UPB820 with chromosomal *xopE5* deletion | This study |
| UPB820 Δ*hopW1* | UPB820 with chromosomal *hopW1* deletion | This study |
| UPB820 Δ*xopAL1/xopAL1* | Complemented strain of UPB820 Δ*xopAL1* | This study |
| UPB820 Δ*xopAJ/xopAJ* | Complemented strain of UPB820 Δ*xopAJ* | This study |
| UPB820 WT + pPneo-GFP | UPB820 WT bearing the pPneo-GFP plasmid for GFP expression, $Km^R$ | This study |
| UPB820 Δ*xopAL1* + pPneo-GFP | UPB820 Δ*xopAL1* bearing the pPneo-GFP plasmid for GFP expression, $Km^R$ | This study |
| UPB820 Δ*xopAL1/xopAL1* + pPneo-GFP | UPB820 Δ*xopAL1/xopAL1* bearing the pPneo-GFP plasmid for GFP expression, $Km^R$ | This study |
| UPB787 WT | Wild-type Xtt strain UPB787 | [1] |
| UPB787 Δ*xopAL1* | UPB787 with chromosomal *xopAL1* deletion | This study |
| XtKm7 WT | Wild-type Xtt strain XtKm7 | [3] |
| XtKm7 Δ*xopAL1* | XtKm7 with chromosomal *xopAL1* deletion | This study |
| ***Xanthomonas translucens* pv. undulosa (Xtu)** | | |
| UPB513 WT | Wild-type Xtu strain UPB513 | [1] |
| UPB513 Δ*hrcT* | UPB513 with chromosomal *hrcT* deletion | This study |
| UPB513 Δ*xopE4* | UPB513 with chromosomal *xopE4* deletion | This study |
| UPB513 miniTn*7*::Gm | UPB513 bearing genomic insertion using the pUC18miniTn*7*T-Gm empty vector, $Gm^R$ | This study |
| UPB513 miniTn*7*::*xopAJ* | UPB513 bearing genomic insertion using the pUC18miniTn*7*T-Gm-*xopAJ* vector for expression of *xopAJ* from Xtt UPB820, $Gm^R$ | This study |
| UPB513 miniTn*7*::*xopAL1* | UPB513 bearing genomic insertion using the pUC18miniTn*7*T-Gm-*xopAL1* vector for expression of *xopAL1* from Xtt UPB820, $Gm^R$ | This study |
| UPB513 miniTn*7*::*xopE5* | UPB513 bearing genomic insertion using the pUC18miniTn*7*T-Gm-*xopE5* vector for expression of *xopE5* from Xtt UPB820, $Gm^R$ | This study |
| UPB513 miniTn*7*::*hopW1* | UPB513 bearing genomic insertion using the pUC18miniTn*7*T-Gm-*hopW1* vector for expression of *hopW1* from Xtt UPB820, $Gm^R$ | This study |
| UPB513 WT + pPneo-GFP | UPB513 WT bearing the pPneo-GFP plasmid for GFP expression, $Km^R$ | [4] |

| | | |
|---|---|---|
| UPB513 + pBBR1-*xopAL1*Xcc8004 | UPB513 bearing the pBBR1-*xopAL1*Xcc8004 plasmid for expression of *xopAL1* from *Xanthomonas campestris* pv. campestris (Xcc) strain 8004, $Km^R$ | This study |
| ***Escherichia coli*** | | |
| DH10B | F– *mcrA* Δ(*mrr-hsdRMS-mcrBC*) φ80*lacZ*ΔM15 Δ*lacX74* *recA1 endA1 araD139* Δ (*ara-leu*)7697 *galU* *galK* λ– *rpsL*($Str^R$) *nupG* | ThermoFisher |
| WM3064 | thrB1004 pro thi rpsL hsdS lacZΔM15 RP4-1360 Δ(araBAD)567 ΔdapA1341::[erm pir] | 5 |
| **Plasmids** | | |
| pK18mobsacB | *sacB* mutagenesis plasmid, $Km^R$ | 6 |
| pUC18miniTn*7*T-Gm | Plasmid used for genomic insertion via miniTn*7*, $Gm^R$ | 7 |
| pTNS3 | Plasmid used for transposition of miniTn*7*, $Ap^R$ | 8 |
| pK18mobsacB::Δ*hrcT*XttUPB820 | *sacB* mutagenesis plasmid for Xtt UPB820 Δ*hrcT* deletion, $Km^R$ | This study |
| pK18mobsacB::Δ*xopAJ*XttUPB820 | *sacB* mutagenesis plasmid for Xtt UPB820 Δ*xopAJ* deletion, $Km^R$ | This study |
| pK18mobsacB::Δ*xopAL1*XttUPB820 | *sacB* mutagenesis plasmid for Xtt UPB820 Δ*xopAL1* deletion, $Km^R$ | This study |
| pK18mobsacB::Δ*xopE5*XttUPB820 | *sacB* mutagenesis plasmid for Xtt UPB820 Δ*xopE5* deletion, $Km^R$ | This study |
| pK18mobsacB::Δ*hopW1*XttUPB820 | *sacB* mutagenesis plasmid for Xtt UPB820 Δ*hopW1* deletion, $Km^R$ | This study |
| pK18mobsacB::*xopAL1*XttUPB820-COMP | *sacB* mutagenesis plasmid designed to complement Xtt UPB820 Δ*xopAL1* by inserting *xopAL1* back into its original locus, $Km^R$ | This study |
| pK18mobsacB::Δ*xopAL1*XttUPB787 | *sacB* mutagenesis plasmid for Xtt UPB886 Δ*xopAL1* deletion, $Km^R$ | This study |
| pK18mobsacB::Δ*xopAL1*XttXtKm7 | *sacB* mutagenesis plasmid for Xtt CIX95 Δ*xopAL1* deletion, $Km^R$ | This study |
| pK18mobsacB::Δ*hrcT*XtuUPB513 | *sacB* mutagenesis plasmid for Xtu UPB513 Δ*hrcT* deletion, $Km^R$ | This study |
| pK18mobsacB::Δ*xopE4*XtuUPB513 | *sacB* mutagenesis plasmid for Xtu UPB513 Δ*xopE4* deletion, $Km^R$ | This study |
| pUC18miniTn*7*T-Gm::*xopAJ*XttUPB820 | Plasmid used for miniTn*7*::*xopAJ* (from Xtt UPB820; including native promoter and terminator regions) genomic insertion, $Gm^R$ | This study |
| pUC18miniTn*7*T-Gm::*xopAL1*XttUPB820 | Plasmid used for miniTn*7*::*xopAL1* (from Xtt UPB820; including native promoter and terminator regions) genomic insertion, $Gm^R$ | This study |
| pUC18miniTn*7*T-Gm::*xopE5*XttUPB820 | Plasmid used for miniTn*7*::*xopE5* (from Xtt UPB820; including native promoter and terminator regions) genomic insertion, $Gm^R$ | This study |
| pUC18miniTn*7*T-Gm::*hopW1*XttUPB820 | Plasmid used for miniTn*7*::*hopW1* (from Xtt UPB820; including native promoter and terminator regions) genomic insertion, $Gm^R$ | This study |

| | | |
|---|---|---|
| pPneo-GFP | Plasmid for GFP expression under the control of the promoter of neomycin resistant gene, $Km^R$ | [9] |
| pBBR1MCS-2 | RK2 broad-host-range cloning vector, $Km^R$ | [10] |
| pEL667 | Domesticated GoldenGate-compatible derivative of pBBR1MCS-2, *Bsa*I cohesive ends GGAG and AGCG, $Km^R$ | This study |
| pEL667-*xopAL1*$_{Xcc8004}$ | pEL667 carrying *xopAL1* from Xcc 8004 including native promoter and terminator regions, $Km^R$ | This study |
| pRK2073 | Helper plasmid, $Sm^R$ | [11] |

$Km^R$ – kanamycin-resistant; $Gm^R$ – gentamicin-resistant; $Rif^R$ – rifampicin-resistant; $Ap^R$ – ampicillin-resistant; $Sm^R$ – streptomycin-resistant.

**Table S7.** List of genes deleted in *X. translucens* in this study.

| Locus tag[a] | Gene | Genome annotation[a] |
|---|---|---|
| ***Xanthomonas translucens* pathovar translucens** | | |
| KFS83_06010 | *hrcT* | Type III secretion system export apparatus subunit SctT |
| KFS83_17905 | *xopAJ* | Zeta toxin family protein |
| KFS83_12860 | *xopAL1* | Type III effector |
| KFS83_00330 | *xopE5* | Type III effector HopX1 |
| KFS83_14440 | *hopW1* | Type III effector |
| ***Xanthomonas translucens* pathovar undulosa** | | |
| F0H33_05575 | *hrcT* | EscT/YscT/HrcT family type III secretion system export apparatus protein |
| F0H33_00965 | *xopE4* | Hypothetical protein |

[a]Locus tags and genome annotations are from genomes available online at NCBI, which are used here as references. Strain UPB947 is the reference for *X. translucens* pv. translucens (GenBank Accession number CP076248), and strain P3 is the reference for *X. translucens* pv. undulosa (GenBank Accession number CP043500).

**Table S8.** Oligonucleotide primers used in this study.

| Name | Sequence (5'-3') | Purpose | Amplicon size (bp) | Source |
|---|---|---|---|---|
| **Oligonucleotide primers used to knockout genes of interest in respective strains** | | | | |
| XttUPB820-hrcT-UP-F | ctctagagtcgacctgcaggcatgcaagctgat gtgaccgatttccagcagtcgctgtgg | Amplify the upstream region of *hrcT* (Xtt UPB820) for cloning into pK18mobsacB | 663 | This study |
| XttUPB820-hrcT-UP-R | agagcgcaactcatacagtgcgcgatggtccat gaacagcacgaagcaacgcacgctcgc | | | This study |
| XttUPB820-hrcT-Down-F | gcgagcgtgcgttgcttcgtgctgttcatggacc atcgcgcactgtatgagttgcgctct | Amplify the downstream region of *hrcT* (Xtt UPB820) for cloning into pK18mobsacB | 549 | This study |
| XttUPB820-hrcT-Down-R | acgttgtaaaacgacggccagtgccaagctctg ctgaagcatctgccaggcggtattacg | | | This study |
| hrcT-conf-F | cttcgtgctgttcatggtgctg | Amplify an internal region of *hrcT* (Xtt UPB820 and Xtu UPB513) to confirm deletion | 679 | This study |
| hrcT-conf-R | gaaggacagttcgtccagcaac | | | This study |
| XttUPB820-xopAJ-UP-F | ctctagagtcgacctgcaggcatgcaagctcat cgtcggatcgctcagcgcctaccattg | Amplify the upstream region of *xopAJ* (Xtt UPB820) for cloning into pK18mobsacB | 533 | This study |
| XttUPB820-xopAJ-UP-R | ctgccaagccatatttttcgctctcaatccccagg aaaatgcgatgacccacttaaaagg | | | This study |
| XttUPB820-xopAJ-Down-F | ccttttaagtgggtcatcgcattttcctggggattg agagcgaaaaatatggcttggcag | Amplify the downstream region of *xopAJ* (Xtt UPB820) for cloning into pK18mobsacB | 537 | This study |
| XttUPB820-xopAJ-Down-R | acgttgtaaaacgacggccagtgccaagctgc cgtagacgtcgccgatgaacaggttgag | | | This study |
| xopAJXttUPB820 conf-F | gatgtttggactgggtgagcttc | Amplify an internal region of *xopAJ* (Xtt UPB820) to confirm deletion | 670 | This study |
| xopAJXttUPB820 conf-R | gaacctgtcgatgacatctaagc | | | This study |
| XttUPB820-xopAL1-UP-F | ctctagagtcgacctgcaggcatgcaagctacc gtatcccgatcgtggcggtgaccaatc | Amplify the upstream region of *xopAL1* (Xtt UPB820) for cloning into pK18mobsacB | 801 | This study |
| XttUPB820-xopAL1-UP-R | cgtacgacactacaacgcctcatgatcgactgc aactgaagagcggcgagcaacccgcag | | | This study |
| XttUPB820-xopAL1-Down-F | ctgcgggttgctcgccgctcttcagttgcagtcg atcatgaggcgttgtagtgtcgtacg | Amplify the downstream region of *xopAL1* (Xtt UPB820) for cloning into pK18mobsacB | 577 | This study |
| XttUPB820-xopAL1-Down-R | acgttgtaaaacgacggccagtgccaagctaa cccgttccacccgttcagcgtgttcgac | | | This study |
| xopAL1Xtt-conf-F | ccttatggcgctggaaatcag | Amplify an internal region of *xopAL1* (all Xtt strains) to confirm | 605 | This study |
| xopAL1Xtt-conf-R | gacatgatcgacgagcgatac | | | This study |

| | | | | |
|---|---|---|---|---|
| | | deletion and complementation | | |
| XttUPB820-xopE5-UP-F | ctctagagtcgacctgcaggcatgcaagctgatcttcaccggcaacgacgaaatggcggc | Amplify the upstream region of *xopE5* (Xtt UPB820) for cloning into pK18mobsacB | 585 | This study |
| XttUPB820-xopE5-UP-R | gcaccaggtgaacgccgtcaggcggggcgcacgcaactccaatgtgtgcaagcgcccgcg | | | This study |
| XttUPB820-xopE5-Down-F | cgcgggcgcttgcacacattggagttgcgtgcgccccgcctgacggcgttcacctggtgc | Amplify the downstream region of *xopE5* (Xtt UPB820) for cloning into pK18mobsacB | 720 | This study |
| XttUPB820-xopE5-Down-R | acgttgtaaaacgacggccagtgccaagctcgatgtactggtagatggtgcgcgcgttgc | | | This study |
| xopE5XttUPB820 conf-F | gttcgtcgagtatgtcgagcaaatg | Amplify an internal region of *xopE5* (Xtt UPB820) to confirm deletion | 503 | This study |
| xopE5XttUPB820 conf-R | catcgttgagcacatgctgttc | | | This study |
| XttUPB820-hopW1-UP-F | ctctagagtcgacctgcaggcatgcaagctaagtctgccaaaaaccgtaagcacatcacc | Amplify the upstream region of *hopW1* (Xtt UPB820) for cloning into pK18mobsacB | 720 | This study |
| XttUPB820-hopW1-UP-R | cggctatgaatctctcgctcataggcggattgaacaaagcatagctcgcggctgagtgta | | | This study |
| XttUPB820-hopW1-Down-F | tacactcagccgcgagctatgctttgttcaatccgcctatgagcgagagattcatagccg | Amplify the downstream region of *hopW1* (Xtt UPB820) for cloning into pK18mobsacB | 632 | This study |
| XttUPB820-hopW1-Down-R | acgttgtaaaacgacggccagtgccaagcttatccgctgcattcggatggcacagcagta | | | This study |
| hopW1XttUPB820conf-F | gcaacaatgctcgatctagatgatgg | Amplify an internal region of *hopW1* (Xtt UPB820) to confirm deletion | 592 | This study |
| hopW1XttUPB820conf-R | ccatcgtggacaaaggtaaaagacg | | | This study |
| XtuUPB513-hrcT-UP-F | ctctagagtcgacctgcaggcatgcaagctgacgatgatgcaggtattgcaacgcctgcg | Amplify the upstream region of *hrcT* (Xtu UPB513) for cloning into pK18mobsacB | 682 | This study |
| XtuUPB513-hrcT-UP-R | gtgcgttgcttcgtgctgttcatggtgctgttgctgatgctggcgttgctgatgcaagtg | | | This study |
| XtuUPB513-hrcT-Down-F | cacttgcatcagcaacgccagcatcagcaacagcaccatgaacagcacgaagcaacgcac | Amplify the downstream region of *hrcT* (Xtu UPB513) for cloning into pK18mobsacB | 719 | This study |
| XtuUPB513-hrcT-Down-R | acgttgtaaaacgacggccagtgccaagctatttccagcagtcgctgtggcacttgcagc | | | This study |
| xopE4XtuUPB513 conf-F | gatgcaatcgccggagctatcatc | Amplify an internal region of *xopE4* (Xtu | 638 | This study |

| | | | | |
|---|---|---|---|---|
| xopE4XtuUPB513 conf-R | ctctgcttttgtccatcgaccaagc | UPB513) to confirm deletion | | This study |
| M13-F | gtaaaacgacggccag | Amplify an internal region of pK18mobsacB to confirm cloning of regions of interest | variable | [12] |
| M13-R | caggaaacagctatgac | | | |
| **Oligonucleotide primers used to insert genes into XtuUPB513 via miniTn*7*** | | | | |
| XttUPB820COMPxopAJ-ORF-F | cttggcgcttttaaaaatttccttggccgccgcatttcctggtcagatcaatgaactgt | Amplify *xopAJ* (Xtt UPB820) including promoter (272 bp) for cloning into pUC18miniTn*7*T-Gm | 1,542 | This study |
| XttUPB820COMPxopAJ-ORF-R | aaggccttcgcgaggtaccgggcccaagctgcgagaggcgccagtcatgtgtgttgcaac | | | This study |
| XttUPB820COMPxopAJ-term-F | ccccgggctgcaggaattcctcgagaagctcatcgtcggatcgctcagcgcctaccattg | Amplify terminator region of *xopAJ* (Xtt UPB820) for cloning into pUC18miniTn*7*T-Gm | 223 | This study |
| XttUPB820COMPxopAJ-term-R | acagttcattgatctgaccaggaaaatgcggcggccaaggaaatttttaaaagcgccaag | | | This study |
| XttUPB820COMPxopAL1-F | ccccgggctgcaggaattcctcgagaagctacgaatagcaggttgcgggtcatgtcgcgt | Amplify *xopAL1* (Xtt UPB820) including promoter (338 bp) and terminator (248 bp) for cloning into pUC18miniTn*7*T-Gm | 1,504 | This study |
| XttUPB820COMPxopAL1-R | aaggccttcgcgaggtaccgggcccaagctccgaaggacaaccgctcatccaagacgacg | | | This study |
| XttUPB820COMPxopE5-F | ccccgggctgcaggaattcctcgagaagctaagactgatccgaggcgagcgtcagcctgc | Amplify *xopE5* (Xtt UPB820) including promoter (313 bp) and terminator (163 bp) for cloning into pUC18miniTn*7*T-Gm | 1,598 | This study |
| XttUPB820COMPxopE5-R | aaggccttcgcgaggtaccgggcccaagctgtacgggccattctagccgccatggttaac | | | This study |
| XttUPB820COMPhopW1-F | ccccgggctgcaggaattcctcgagaagctcctctcaacatacaccggtgctcacatcaa | Amplify *hopW1* (Xtt UPB820) including promoter (405 bp) and terminator (265 bp) for cloning into pUC18miniTn*7*T-Gm | 1,960 | This study |
| XttUPB820COMPhopW1-R | aaggccttcgcgaggtaccgggcccaagctcagaacacgtgatcgatcaaatggtcgcgc | | | This study |
| miniTN7-conf-F | cgttcggtcaaggttctgga | Amplify an internal region of pUC18miniTn*7*T-Gm to confirm cloning of regions of interest | variable | This study |
| miniTn7-conf-R | gctggccgataagctctgat | | | This study |
| **Oligonucleotide primers used to clone *xopAL1*$_{\text{Xcc8004}}$ into pEL667 replicative plasmid** | | | | |
| LM-169-XC_2995 xopAL1 | tggtctctggagttctgaagcatgtggtcgg | Amplify *xopAL1* 5' region for Golden Gate cloning into pEL667 | 850 | This study |
| LM170-XC_2995 xopAL1 | tggtctcgtcctcatctgcgtcgcttactc | | | |
| LM171-XC_2995 xopAL1 | tggtctcgaggaccatgcttacgtgctga | | 568 | This study |

| | | |
|---|---|---|
| LM-172-XC_2995 xopAL1 | tggtctcaagcgttcctcggagtaggaaatca | Amplify *xopAL1* 3’ region for Golden Gate cloning into pEL667 |

**Table S9.** List of genomes used in this study.

| Strain | GenBank Accession number | Reference |
|---|---|---|
| ***Xanthomonas translucens* pv. translucens (Xtt)** | | |
| B1FA | CP090000 | Unpublished |
| CIX43 | CP072988/CP072989 | [13] |
| CIX84 | CP159782 | This study |
| CIX95 | CP072990 | [13] |
| Colorado | https://github.com/Merfa-lab/Genomes | This study |
| DSM 18974 | LT604072 | [14] |
| UPB458 | CP076249 | [15] |
| UPB545 | CP159781 | This study |
| UPB787 | https://github.com/Merfa-lab/Genomes | This study |
| UPB820[a] | https://github.com/Merfa-lab/Genomes | This study |
| UPB886 | GCA_009600865.1 | [16] |
| UPB947 | CP076248 | [15] |
| XtKm7 | CP064005 | [3] |
| XtKm8 | CP064004 | [3] |
| XtKm9 | CP064003 | [3] |
| XtKm34 | CP064001 | [3] |
| ***Xanthomonas translucens* pv. undulosa (Xtu)** | | |
| CFBP 2055 | CP074361/CP074362 | [15] |
| CFBP 2539 | CP074363 | [15] |
| CIX162 | CP093449 | [17] |
| CIX207 | CP093448 | [17] |
| CIX282 | CP093447 | [17] |
| CIX303 | CP093446 | [17] |
| CIX40 | CP093450 | [17] |
| ICMP 11055 | CP009750 | [18] |
| LW16 | CP043540 | [19] |
| MAI5034 | CP089584 | [20] |
| P3 | CP043500 | [19] |
| UPB513[a] | JBEFAI000000000 | This study |
| XtFa1 | CP063996 | [3] |
| XtKm12 | CP064000 | [3] |
| XtKm15 | CP063997/CP063998/CP063999 | [3] |
| XtLr8 | CP063993/CP063994/CP063995 | [3] |
| Xtu 4699 | CP008714 | Unpublished |
| ***Xanthomonas translucens* pv. arrhenateri (Xta)** | | |
| LMG 727 | CP086333 | [15] |
| ***Xanthomonas translucens* pv. cerealis (Xtc)** | | |
| 01 | CP038228 | [21] |
| CFBP 2541 | CP074364 | [15] |
| ***Xanthomonas translucens* pv. graminis (Xtg)** | | |
| LMG 726 | CP076254 | [15] |
| ART-Xtg9 | CP076252 | [22] |
| ART-Xtg2 | CP076253 | [22] |
| ART-Xtg29 | CP076257 | [22] |

| | | |
|---|---|---|
| NCPPB 3709 | CP118161/CP118162 | [22] |
| ***Xanthomonas translucens* pv. phleipratensis (Xtphlei)** | | |
| LMG 730 | CP076251 | [15] |
| LMG 843 | CP086332 | [15] |
| ***Xanthomonas translucens* pv. pistaciae (Xtp)** | | |
| CFBP 8304 | CP074365 | [15] |
| ICMP 16317 | CP083804 | [15] |
| ***Xanthomonas translucens* pv. poae (Xtpo)** | | |
| LMG 728 | CP076250 | [15] |
| NCPPB 3711 | CP076255/CP076256 | Unpublished |
| ***Xanthomonas albilineans*** | | |
| Xa-FJ1[a] | CP046570.1/CP046571.1 | [23] |
| ***Xanthomonas arboricola* (Xa)** | | |
| YchA | CP090441 | Unpublished |
| ***Xanthomonas arboricola* pv. juglandis** | | |
| CFBP 2528[a] | GCA_001013475.1 | [24] |
| ***Xanthomonas axonopodis* pv. cassiae** | | |
| NCPPB 645 | CP168000.1 | Unpublished |
| ***Xanthomonas axonopodis* pv. vasculorum** | | |
| NCPPB 796[a] | CP053649.1/CP053650.1/CP053651.1 | Unpublished |
| ***Xanthomonas bromi*** | | |
| LMG 947 | GCA_900092025.1 | Unpublished |
| ***Xanthomonas campestris* (Xc) pv. campestris (Xcc)** | | |
| 40-2 | CP067001 | [25] |
| 3054 | CP066963/CP066964/CP066965 | [25] |
| 8004 | CP000050 | [26] |
| CFBP 12825 | CP066929 | [25] |
| CFBP 5251[a] | AE008922.1 | [27] |
| CN07 | CP101174/CP101175 | Unpublished |
| ***Xanthomonas cannabis*** | | |
| 8590[a] | GCA_011761725.1 | Unpublished |
| ***Xanthomonas euvesicatoria* (Xe)** | | |
| T0319-01 | CP137539.1 | [28] |
| ***Xanthomonas euvesicatoria* pv. perforans** | | |
| GEV872[a] | CP116305.1/CP116306.1/CP116307.1/CP116308.1 | Unpublished |
| ***Xanthomonas euvesicatoria* pv. polysciadis** | | |
| Pgu1 | CP180391.1 | Unpublished |
| ***Xanthomonas fragariae* (Xf)** | | |
| YLX21 | CP114134.1 | [29] |

| | | |
|---|---|---|
| ***Xanthomonas hortorum*** | | |
| B07-007[a] | CP016878.1/CP016879.1 | Unpublished |
| ***Xanthomonas hortorum* pv. hederae (Xhh)** | | |
| 22-338 | GCA_028580375.1 | [30] |
| ***Xanthomonas hortorum* pv. pelargonii (Xhp)** | | |
| OSU493 | CP098604/CP098605 | Unpublished |
| OSU498 | CP098602/CP098603 | Unpublished |
| ***Xanthomonas hortorum* pv. vitians (Xhv)** | | |
| LM16734 | CP060399.1 | [31] |
| ***Xanthomonas hyacinthi*** | | |
| CFBP 1156 | CP043476/CP043477 | [32] |
| ***Xanthomonas nasturtii* (Xn)** | | |
| 10015A | GCA_023337265.1 | Unpublished |
| ***Xanthomonas oryzae* pv. oryzicola** | | |
| GX01[a] | CP043403.1/KR071788.1 | Unpublished |
| BLS256 | CP003057.2 | [33] |
| ***Xanthomonas populi* (Xp)** | | |
| CFBP 1817 | GCA_002940065.1 | Unpublished |
| ***Erwinia psidii* (Ep)** | | |
| LPF 534 | GCA_026549045.1 | [34] |
| ***Paraburkholderia nemoris* (Pn)** | | |
| RL18-011-BIC-A | GCA_046027695.1 | [35] |
| ***Paracidovorax citrulli*** | | |
| AAC00-1 | CP000512.1 | Unpublished |
| DSM 17060 | GCA_900100305.1 | Unpublished |
| ***Pseudomonas amygdali* (Pa)** | | |
| CFBP 1650 | GCA_041449455.1 | Unpublished |
| ***Ralstonia mannitolilytica* (Rm)** | | |
| LMG 18090 | GCA_958405575.1 | [36] |
| ***Ralstonia solanacearum*** | | |
| 10314 | GCA_008271875.1 | [37] |

[a] Indicates genomes that were used for representative *Xanthomonas* spp. phylogenomics and host range in figures 1A and S1.

**Supplementary Materials References**

1 Bragard, C. *et al.* Xanthomonas translucens from small grains: diversity and phytopathological relevance. *Phytopathology* **87**, 1111–1117 (1997).
2 Alizadeh, A., Barrault, G., Sarrafi, A., Rahimian, H. & Albertini, L. Identification of bacterial leaf streak of cereals by their phenotypic characteristics and host range in Iran. *European Journal of Plant Pathology* **101**, 225–229 (1995).
3 Shah, S. M. A. *et al.* Genomics-enabled novel insight into the pathovar-specific population structure of the bacterial leaf streak pathogen Xanthomonas translucens in small grain cereals. *Front Microbiol* **12**, 674952 (2021). https://doi.org/10.3389/fmicb.2021.674952
4 Gluck-Thaler, E. *et al.* Repeated gain and loss of a single gene modulates the evolution of vascular plant pathogen lifestyles. *Sci Adv* **6**, eabc4516 (2020).
5 Dehio, C. & Meyer, M. Maintenance of broad-host-range incompatibility group P and group Q plasmids and transposition of Tn5 in Bartonella henselae following conjugal plasmid transfer from Escherichia coli. *J Bacteriol* **179**, 538–540 (1997).
6 Schäfer, A. *et al.* Small mobilizable multi-purpose cloning vectors derived from the Escherichia coli plasmids pK18 and pK19: selection of defined deletions in the chromosome of Corynebacterium glutamicum. *Gene* **145**, 69–73 (1994).
7 Choi, K. H. & Schweizer, H. P. mini-Tn7 insertion in bacteria with single attTn7 sites: example Pseudomonas aeruginosa. *Nat Protoc* **1**, 153–161 (2006). https://doi.org/10.1038/nprot.2006.24
8 Choi, K. H. *et al.* Genetic tools for select-agent-compliant manipulation of Burkholderia pseudomallei. *Appl Environ Microbiol* **74**, 1064–1075 (2008). https://doi.org/10.1128/aem.02430-07
9 Han, S. W., Park, C. J., Lee, S. W. & Ronald, P. C. An efficient method for visualization and growth of fluorescent Xanthomonas oryzae pv. oryzae in planta. *BMC Microbiol* **8**, 164 (2008). https://doi.org/10.1186/1471-2180-8-164
10 Kovach, M. E. *et al.* Four new derivatives of the broad-host-range cloning vector pBBR1MCS, carrying different antibiotic-resistance cassettes. *Gene* **166**, 175–176 (1995).
11 Leong, S. A., Ditta, G. S. & Helinski, D. R. Heme biosynthesis in Rhizobium: identification of a cloned gene coding for δ-aminolevulininc acid synthetase from Rhizobium meliloti. *J. Biol. Chem.* **257**, 8724–8730 (1982). https://doi.org/10.1016/s0021-9258(18)34188-7
12 Messing, J. New M13 vectors for cloning. *Methods Enzymol* **101**, 20–78 (1983).
13 Heiden, N., Roman-Reyna, V., Curland, R. D., Dill-Macky, R. & Jacobs, J. M. Comparative genomics of barley-infecting Xanthomonas translucens shows overall genetic similarity but globally distributed virulence factor diversity. *Phytopathology* **113**, 2056–2061 (2023). https://doi.org/10.1094/phyto-04-22-0113-sc
14 Jaenicke, S. *et al.* Complete genome sequence of the barley pathogen Xanthomonas translucens pv. translucens DSM 18974T (ATCC 19319T). *Genome Announc* **4** (2016). https://doi.org/10.1128/genomeA.01334-16
15 Goettelmann, F. *et al.* Complete genome assemblies of all Xanthomonas translucens pathotype strains reveal three genetically distinct clades. *Front Microbiol* **12**, 817815 (2021). https://doi.org/10.3389/fmicb.2021.817815

16 Roman-Reyna, V. *et al.* Genome resource of barley bacterial blight and leaf streak pathogen Xanthomonas translucens pv. translucens strain UPB886. *Plant Dis* **104**, 13–15 (2020). https://doi.org/10.1094/pdis-05-19-1103-a
17 Ledman, K. E. *et al.* Comparative genomics of Xanthomonas translucens pv. undulosa strains isolated from weedy grasses and cultivated wild rice. *Phytopathology* **113**, 2083–2090 (2023).
18 Falahi Charkhabi, N. *et al.* Complete genome sequencing and targeted mutagenesis reveal virulence contributions of Tal2 and Tal4b of Xanthomonas translucens pv. undulosa ICMP11055 in bacterial leaf streak of wheat. *Front Microbiol* **8**, 1488 (2017). https://doi.org/10.3389/fmicb.2017.01488
19 Peng, Z. *et al.* Xanthomonas translucens commandeers the host rate-limiting step in ABA biosynthesis for disease susceptibility. *Proc Natl Acad Sci U S A* **116**, 20938–20946 (2019). https://doi.org/10.1073/pnas.1911660116
20 Clavijo, F. *et al.* Complete genome sequence resource for Xanthomonas translucens pv. undulosa MAI5034, a wheat pathogen from Uruguay. *Phytopathology* **112**, 2036–2039 (2022). https://doi.org/10.1094/phyto-01-22-0025-a
21 Shah, S. M. A. *et al.* Tal1(NXtc01) in Xanthomonas translucens pv. cerealis contributes to virulence in bacterial leaf streak of wheat. *Front Microbiol* **10**, 2040 (2019). https://doi.org/10.3389/fmicb.2019.02040
22 Goettelmann, F., Koebnik, R., Roman-Reyna, V., Studer, B. & Kolliker, R. High genomic plasticity and unique features of Xanthomonas translucens pv. graminis revealed through comparative analysis of complete genome sequences. *BMC Genomics* **24**, 741 (2023). https://doi.org/10.1186/s12864-023-09855-8
23 Zhang, H. L. *et al.* Complete genome sequence reveals evolutionary and comparative genomic features of Xanthomonas albilineans causing sugarcane leaf scald. *Microorganisms* **8** (2020). https://doi.org/10.3390/microorganisms8020182
24 Cesbron, S. *et al.* Comparative genomics of pathogenic and nonpathogenic strains of Xanthomonas arboricola unveil molecular and evolutionary events linked to pathoadaptation. *Front Plant Sci* **6**, 1126 (2015). https://doi.org/10.3389/fpls.2015.01126
25 Dubrow, Z. E. *et al.* Cruciferous weed isolates of Xanthomonas campestris yield insight into pathovar genomic relationships and genetic determinants of host and tissue specificity. *Mol Plant Microbe Interact* **35**, 791–802 (2022). https://doi.org/10.1094/mpmi-01-22-0024-r
26 Qian, W. *et al.* Comparative and functional genomic analyses of the pathogenicity of phytopathogen Xanthomonas campestris pv. campestris. *Genome Res* **15**, 757–767 (2005). https://doi.org/10.1101/gr.3378705
27 da Silva, A. C. R. *et al.* Comparison of the genomes of two Xanthomonas pathogens with differing host specificities. *Nature* **417**, 459–463 (2002). https://doi.org/10.1038/417459a
28 Huang, C. J., Wu, T. L., Wu, Y. L., Wang, R. S. & Lin, Y. C. Comparative genomic analysis uncovered phylogenetic diversity, evolution of virulence factors, and horizontal gene transfer events in tomato bacterial spot Xanthomonas euvesicatoria. *Front Microbiol* **15**, 1487917 (2024). https://doi.org/10.3389/fmicb.2024.1487917
29 Wei, F. *et al.* Pan-Genomic Analysis Identifies the Chinese Strain as a New Subspecies of Xanthomonas fragariae. *Plant Disease* **108**, 45–49 (2023). https://doi.org/10.1094/PDIS-05-23-0933-SC

30 Iruegas-Bocardo, F. *et al.* Whole genome sequencing-based tracing of a 2022 introduction and outbreak of Xanthomonas hortorum pv. pelargonii. *Phytopathology* **113**, 975–984 (2023). https://doi.org/10.1094/phyto-09-22-0321-r

31 Morinière, L., Lecomte, S., Gueguen, E. & Bertolla, F. In vitro exploration of the Xanthomonas hortorum pv. vitians genome using transposon insertion sequencing and comparative genomics to discriminate between core and contextual essential genes. *Microb Genom* **7** (2019). https://doi.org/10.1099/mgen.0.000546

32 Cohen, S. P. *et al.* High-quality genome resource of Xanthomonas hyacinthi generated via long-read sequencing. *Plant Dis* **104**, 1011–1012 (2020). https://doi.org/10.1094/pdis-11-19-2393-a

33 Bogdanove, A. J. *et al.* Two new complete genome sequences offer insight into host and tissue specificity of plant pathogenic Xanthomonas spp. *J Bacteriol* **193**, 5450–5464 (2011). https://doi.org/10.1128/jb.05262-11

34 Alves, F. H. N. d. S. *et al.* Comprehensive prediction of plant cytoplasmic and apoplastic effectors underlying Erwinia psidii pathogenicity. *Plant Pathology* **72**, 130–143 (2023). https://doi.org/https://doi.org/10.1111/ppa.13636

35 Paulo, B. S. *et al.* Discovery of megapolipeptins by genome mining of a Burkholderiales bacteria collection. *Chem Sci* **15**, 16567–16581 (2024). https://doi.org/10.1039/d4sc03594a

36 Steyaert, S. *et al.* Novel Ralstonia species from human infections: improved matrix-assisted laser desorption/ionization time-of-flight mass spectrometry-based identification and analysis of antimicrobial resistance patterns. *Microbiol Spectr* **12**, e0402123 (2024). https://doi.org/10.1128/spectrum.04021-23

37 Bautista, M. A. M., Llames, J. H. S., Sabban, E. A. V. & Villegas, L. C. Draft genome sequences of Ralstonia solanacearum isolated from banana and tomato in the Philippines. *Philippine Journal of Science* **148**, 115–126 (2019).
